# Systematic screen of RNA binding proteins that enhance circular RNA translation

**DOI:** 10.1101/2024.11.07.622558

**Authors:** Qianyun Lu, Siqi Wang, Yanwen Ye, Yun Yang, Zefeng Wang

**Affiliations:** CAS Key Laboratory of Computational Biology, Shanghai Institute of Nutrition and Health, Shanghai Institutes for Biological Sciences; University of Chinese Academy of Sciences, Chinese Academy of Sciences, Shanghai 200031, China; Southern University of Science and Technology, Shenzhen, China; CirCode Biomedicine Inc. Shanghai, China

## Abstract

Translatable circular RNAs (circRNAs) have emerged as a promising alternative to linear mRNA as new therapeutics due to its improved stability. The translation of circRNAs is mainly driving by internal ribosome entry site (IRES) or IRES-like elements, which is under regulation by various *trans*-acting RNA binding proteins (RBPs). Here we designed a cell-based system to systematically screen RBPs that enhance translation of circRNAs, and identified a total of 68 proteins as putative activators of noncanonical translation. These translation activators mainly involved in the functions of RNA processing, ribosomal biogenesis and translation initiation. Furthermore, we developed a machine learning algorithm to extract common sequence features of these activators, which predicted more potential RBPs with translation activator activities. The newly identified and predicted activators were subsequently demonstrated to promote the IRES-mediated circRNA translation in a context-dependent fashion. This investigation provides new insights to discover functions for IRES *trans*-acting factors and to expand the toolbox for engineered RBPs in RNA synthetic biology.

## Introduction

CircRNAs are a class of covalently closed single-stranded RNAs generated in cells mainly through back-splicing. Although discovered more than 30 years ago, only recently people have realized that the circRNAs are abundant and evolutionary conserved among eukaryotes with cell-specific expression patterns (Kristensen et al., 2019; Patop et al., 2019). The circRNAs were thought to function as noncoding RNAs by regulating different steps of gene expression, including transcription, splicing, mRNA stability, translation and signaling pathways through interactions with their target DNAs, RNAs and proteins (Conn et al., 2024; Kristensen et al., 2022; Liu and Chen, 2022). However, certain circRNAs can also be translated into functional proteins (Abe et al., 2015; Chen et al., 2021a; Chen and Sarnow, 1995; Pamudurti et al., 2017; Wang and Wang, 2015; Yang et al., 2017), showing promise as a new type of mRNA vaccine due to their higher cellular stability (Perenkov et al., 2023; Wei et al., 2023). In addition to the superior stability, the properly purified circRNAs have lower immunogenicity even without base modification (Petkovic and Müller, 2015; Wesselhoeft et al., 2018; Wesselhoeft et al., 2019), thus may provide a wider expression window for therapeutic proteins as the next generation of RNA therapy (Holdt et al., 2018; Niu et al., 2023).

Canonical translation of linear mRNA is initiated with the recognition of m7G cap structure by eukaryotic initiation factor 4E (eIF4E), which is in synergy with eukaryotic initiation factor 4G (eIF4G) and 4A (eIF4A) to form the eIF4F complex, followed by the recruitment of other translation initiation factors and the ribosome (Hinnebusch, 2014). When cells undergo stress (e.g., hypoxia, apoptosis, starvation, and viral infection), the canonical protein synthesis is downregulated, while the proteins essential for cell survival and stress recovery are selectively translated (Kwan and Thompson, 2019). A major mechanism involved in this noncanonical translation is through the use of IRES, which are often assisted by certain RBPs known as IRES *trans*-<u>a</u>cting factors (ITAFs), in a cap-independent manner (Godet et al., 2019). The IRES-mediated translation was also observed in naturally occurring (Legnini et al., 2017; Pamudurti et al., 2017; Yang et al., 2018) and synthetically designed circRNAs (Chen et al., 2021a; Chen and Sarnow, 1995; Chen et al., 2023; Fan et al., 2022; Wang and Wang, 2015). However, the efficiency of IRES-driven translation is generally low (Feng et al., 2023), posing a challenge for leveraging circRNAs as therapeutic tools. In addition, the activity of IRES is highly dependent on the expression and activity of specific ITAFs, resulting in cell-type-specific regulation of IRES-mediated translation (Godet et al., 2019). Despite their potential in biomedical applications, the *trans*-acting factors that enhance cap-independent circRNA translation (*i.e.*, functional equivalent to ITAFs) have yet to be thoroughly dissected (Deviatkin et al., 2023). Identifying these factors is critical to improving the efficiency and specificity of circRNA translation, which could pave the way for the development of novel translation manipulation tools and therapeutic applications involving circRNAs.

To study *trans*-acting factors that regulate circRNA translation, we employed a luciferase-based circRNA reporter system coupled with a tethered RBP library to systematically identify putative proteins that can promote circRNA translation. Remarkably, we found that a large number of RBPs tested could facilitate noncanonical translation when binding to the circRNAs. For simplicity, we termed these *trans*-acting RBPs that can enhance circRNA translation as the translation activators in this paper. We further developed a machine learning (ML) model using the experimentally tested proteins as the training dataset to predict additional translation activators from all known RBPs. Both experimentally identified translation activators and predicted activators were shown to stimulate IRES-mediated circRNA translation, suggesting they may function as novel ITAFs for certain IRES. Overall, our study extended the scope of current knowledge on translation regulatory RBPs and uncovered an inherent connection between RBP sequence and their translation promoting activity, which may hold promise for applications in RNA-based therapeutics.

## Results

### Engineer a reporter system to examine RBP activities in regulating circRNA translation

We have previously developed a circEGFP reporter system to identify regulatory *cis*-elements that drive cap-dependent translation of circRNAs (Wang and Wang, 2015; Yang et al., 2017; Yang and Wang, 2019). Building upon a similar design principle, we engineered a luminescence-based system that allows specific detection of translation products from circRNAs *vs.* linear mRNAs. In this system, the coding sequence (CDS) of *firefly* luciferase (Fluc) was split into two fragments in reverse order flanked by two introns with complementary elements (Figure 1A), generating a circular RNA with the intact open reading frame (ORF) of *firefly* luciferase (cFluc) through back-splicing. As an internal control, we incorporated the renilla luciferase (Rluc) gene in an inverse orientation (Figure 1A), which can be used to calibrate for effect from cell transfection and gene expression from linear mRNA.

**Figure 1.**
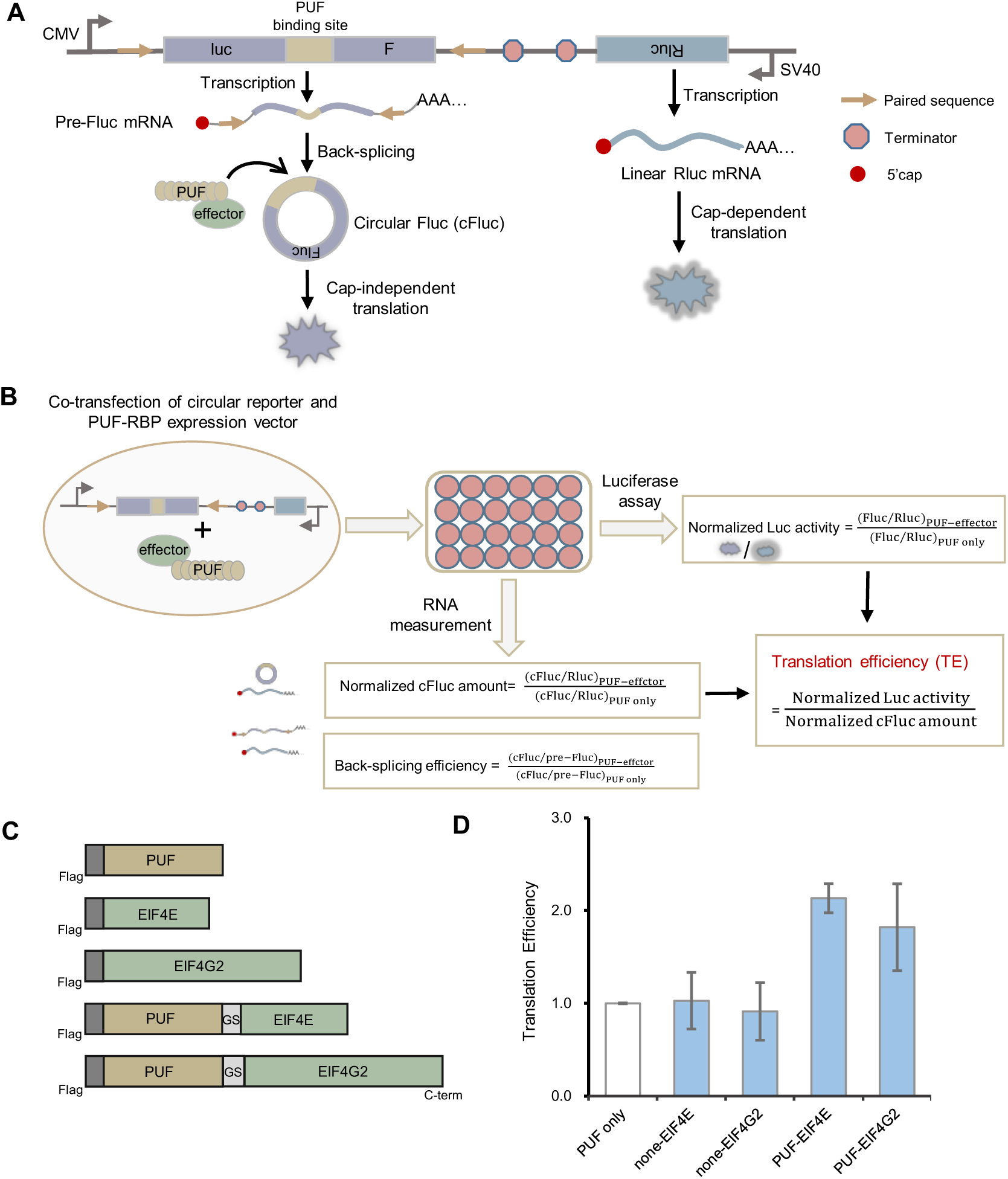
Design and validation of circular reporter system to examine RBP activities. (A) Schematic illustration of the circular luciferase reporter system. The Fluc gene was split into two fragments in reverse order and constructed in a single exon, which can be back-spliced into a circular RNA *via* the complementary sequences in the flanking introns. Two tandem PUF binding sites were inserted between the start and stop codon of cFluc to tether the artificial PUF factors. An Rluc gene was inserted into the same vector as a transfection control. (B) Workflow of the experimental and data analysis. The luciferase activity and RNA levels were measured from the same sample for the back-splicing efficiency and translation efficiency. The measurements were normalized to the values obtained from the PUF only control. (C) The schematic diagram of the constructs for validation: PUF only, none-EIF4E, none-EIF4G2, PUF-EIF4E, PUF-EIF4G2. The PUF domain was located at the N-terminal and the effector domain at the C-terminal. The Flag epitope tag was depicted in dark gray, and the GS linker in light gray. (D) Validation of the reporter system by tethering two translation initiation factors, eIF4E and eIF4G2. Translation efficiency of PUF only, none-EIF4E, none-EIF4G2, PUF-EIF4E, and PUF-EIF4G2 on the cFluc reporter was determined using the luciferase assay described in panel B (mean ± SD, in triplicate transfections).

Next, we used a modified Pumilio and FBF homology (PUF) domain that specifically bind an 8-nucleotide (nt) RNA sequences (UUGAUAUA) to tether candidate RBPs onto the circRNA. The PUF-RBPs were co-transfected with the circular RNA reporter in human embryonic kidney 293T (HEK 293T) cells and analyzed at 36h post transfection (methods). As a control, we utilized a PUF domain fused only with a FLAG tag (PUF-only). We quantified the luciferase activities and RNA levels of cFluc and Ruc, and normalized to the samples co-expressed with the PUF-only control (Figure 1B).

To validate this system, we initially tested two known translation initiation factors, eIF4E and eIF4G2. eIF4E is known to recognize the 5’ m7G cap of mRNA and, in synergy with eIF4A and eIF4G, recruits other initiation factors to activate translation (Hinnebusch, 2014). The eIF4G2 has been reported to facilitate IRES-dependent translation of mRNA (Marash et al., 2008) and the m6A-driven translation of circRNAs (Yang et al., 2017). We fused the ORFs of eIF4E and eIF4G2 with the FLAG epitope and the PUF domain at the N terminus (Figure 1C). As expected, tethering eIF4E and eIF4G2 to the upstream of circRNA start codon both enhanced the translation efficiency of circRNA (Figure 1D). The eIF4E/eIF4G2 fused with FLAG tag didn’t activate the reporter, suggesting that the direct tethering through PUF is essential for their activity. Although eIF4E has not been reported to promote circRNA translation, our findings suggest that the recruitment of eIF4E by an artificial system can promote circRNA translation, probably *via* the association of eIF4E with the translation initiation apparatus. Collectively, this tethering system may serve as both a discovery tool to identify *trans-*acting factors that promote circRNA translation, as well as a platform for new molecular toolkits to optimize circRNA therapy.

### Systematic screen for RBPs that affect circRNA translation

Previous study demonstrated that a large number of ribosome-associated proteins, mostly RBPs, were functionally enriched for IRES-mediated translation and ribosome assembly process (Simsek et al., 2017). We hypothesized that some of these proteins may have the potential to modulate cap-independent circRNA translation. In total 110 putative RBPs were selected into the candidate library and the full-length CDS of these proteins were cloned into the PUF constructs (Figure 2A) (methods). Gene Ontology (GO) analysis of 110 RBPs revealed a significant enrichment in the core pathways related to RNA splicing, translation initiation and ribosomal biogenesis (Supplementary Figure S1A). Moreover, most of these RBPs are highly expressed as judged by RNA-seq and ribosome profiling data (Calviello et al., 2016) (Supplementary Figure S1B), consistent with a general role in mediating housekeeping pathways of cells.

**Figure 2.**
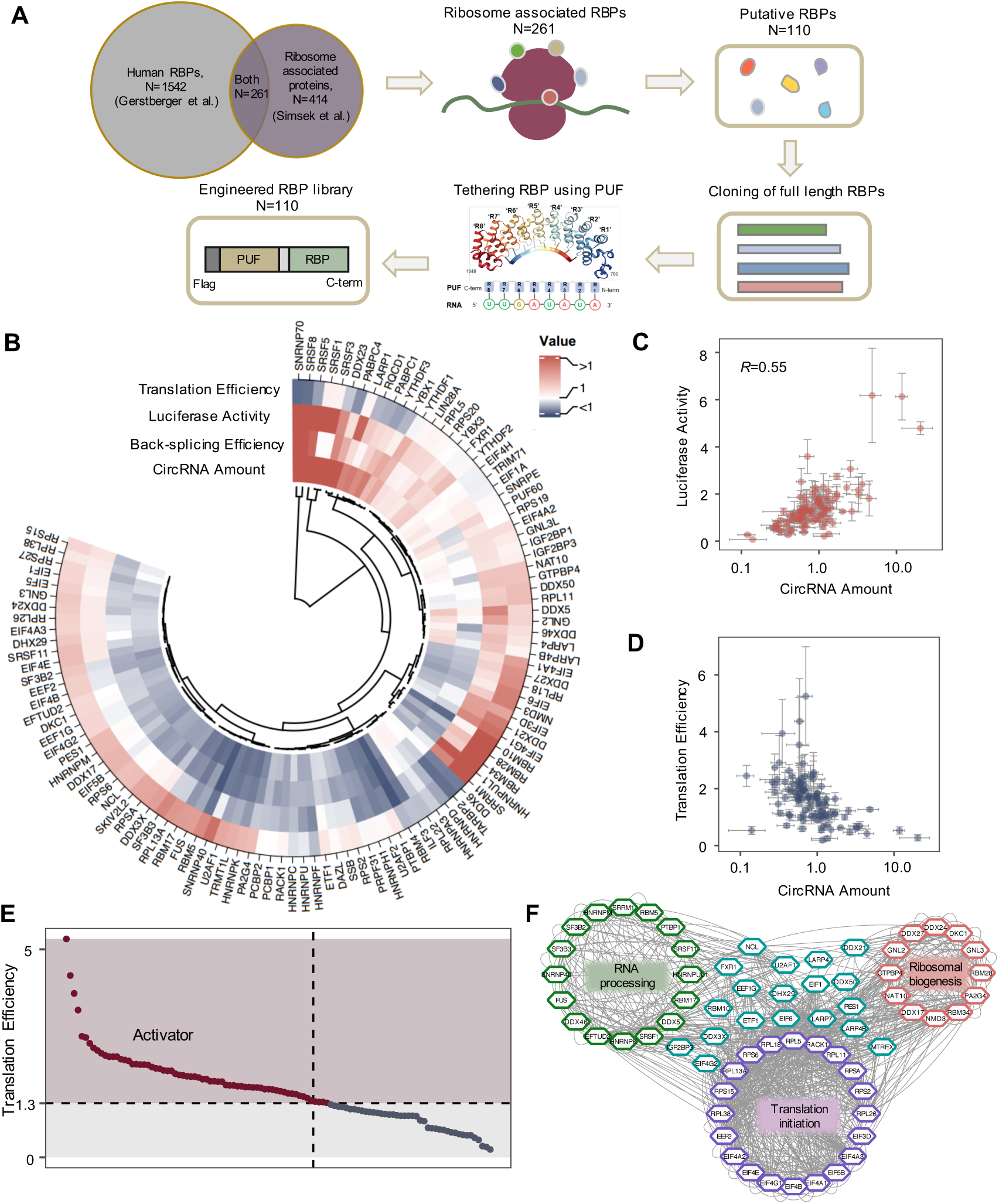
Screen for RBPs that promote circRNA translation. (A) Schematic diagram for selection of the 110 putative RBPs and construction of the PUF-RBP fusion protein library. (B) Hierarchical clustering (with Ward’s minimum variance method) of luciferase activity, circRNA amount, back splicing efficiency and translation efficiency (TE) for 110 RBPs as calculated from the workflow in Fig 1(B) (mean ± SD, in triplicate transfections). (C) The correlation between luciferase activity and circRNA amount for all the 110 RBPs tested (*R*, Pearson correlation coefficient). (D) The correlation between translation efficiency and circRNA amount for the 110 RBPs. (E) Scatter plot showing the distribution of TE values for all tested RBPs, where the RBPs with increased TE value (TE ≥ 1.3) were identified as translation activators. (F) Functional network of the identified activators. The activators were analyzed using STRING database and clustered into three main groups by CytoScape MCODE tool.

Next, we transfected all tethering RBPs in triplicate and compared them with the PUF-only control transfected on the same plate. Protein expression levels were validated through western blotting analysis of cell lysates (Supplementary Figure S1C). When co-expressed with the dual luciferase reporter, PUF-RBPs may affect cFluc signals by regulating back-splicing efficiency and/or cap-independent translation. Therefore we measured the RNA levels of both linear and circular mRNAs, as well as the activities of both luciferases, to calculate their effect on both translation and back-splicing (Figure 1B). Subsequently, the translation efficiency, luciferase activity, splicing efficiency and circRNA amount affected by all 110 RBPs were clustered and plotted using a heatmap (Figure 2B). We observed a general positive correlation between luciferase activity and circRNA amount (*R* = 0.55) (Figure 2C), while TE exhibited an inverse correlation with circRNA amount (Figure 2D). Additionally, a significant subset of RBPs was found to reduce circRNA splicing, leading to a decrease in circRNA levels (Supplementary Figure S2A, B). These findings were consistent with the notion that back-splicing may affect translation efficiency by altering circRNA abundance.

### Characteristics for identified RBPs that activate circRNA translation

We further focused on RBPs that increased relative translation efficiency (TE) of circRNAs by at least 30% (*i.e.*, using relative TE cutoff of 1.3). Of the 110 RBPs examined, 68 were identified as the translation activators (Figure 2E). We first investigated the underlying biological functions of these candidate activators using GO analysis, and found strong functional enrichments in translation initiation (e.g., eukaryotic initiation factors (eIFs) and ribosomal proteins), RNA processing (e.g., hnRNPs) and ribosomal biogenesis (e.g., DDX family) (Figure 2F). These enrichments were consistent with our screening design and objectives, which further reassured the results. Notably, many identified translation activators, such as hnRNPUL1, RPS2, IGF2BP3 and SRSF1, were previously reported to bind IRES-like short elements in a circular GFP system (Fan et al., 2022). In addition, hnRNPM was reported as an ITAF to induce translational switch under hypoxia to promote colon cancer development (Chen et al., 2019). Interestingly, we identified more than 10 members from DEAD-box helicase family as translation activators (Figure 2F), some of which had been previously shown to possess ITAF activities. For example, DDX3X was documented to engage with a number of 40S ribosomal components and facilitate IRES-driven translation in FMDV (Han et al., 2020). Additionally, the DEAD-box helicase component of the exon-junction complex, eIF4A3, was recently reported to promote circRNA translation *via* interaction with eIF3G (Chang et al., 2023).

To examine whether these identified activators of cap-independent translation exhibit distinct functions in linear *vs.* circular mRNAs, we utilized a standard bicistronic reporter (Carter and Sarnow, 2000). We introduced two consecutive PUF binding motifs between the start codon of linear Fluc and the stop codon of linear Rluc in the reporter construct (Supplementary Figure S3A). A total of 15 tethered RBPs with a range of TE values, were tested by co-transfecting with the reporter into HEK 293T cells. The results revealed a strong positive correlation between the translation enhancing activities of RBPs measured by this bicistronic reporter system and their activities observed in the circRNA context (Supplementary Figure S3B), suggesting potential general roles of certain RBPs in promoting cap-independent translation.

Given the significance of RBP intracellular localization in modulating RNA metabolism (Benoit Bouvrette et al., 2022), we clustered our translation activators based on their subcellular localization information from Uniprot database. This analysis uncovered that the majority of the 68 translation activators located in both nucleus and cytoplasm (Supplementary Figure S3C), consistent with previous finding that most known ITAFs regulate IRES-dependent translation by shuttling between the two cellular compartments (Godet et al., 2019). Interestingly, of the 68 translation activators, 27 were associated with stress granules that contain many specific translation initiation factors, while 12 were mapped to P bodies that are enriched with factors in mRNA degradation and decay (Riggs et al., 2020). This distribution pattern implies that these RBPs may play an ITAF-like role in regulating circRNA translation under cellular stress.

### Exploring endogenous roles for the newly identified translation activators

To our knowledge, only a limited number of *trans-*factors were reported to drive circRNA translation (Chang et al., 2023; Fan et al., 2022). On the other hand, approximately 55 ITAFs have been documented to modulate cellular IRESs under different physiological conditions, predominantly functioning as activators (Godet et al., 2019). In light of this, we compared our identified activators with the 55 reported cellular ITAFs (Supplementary Figure S4A). Our analysis revealed that 14 out of the 68 identified activators were previously reported as ITAFs (eIF4A1, eIF4G1, eIF4G2, eIF5B, FUS, hnRNPK, hnRNPM, NCL, PA2G4, PTBP1, RACK1, RPL11, RPL26, RPL38), while 54 novel translation activators were identified (Fig 3A). Notably, many members from the La-related protein family (LARP1, LARP4, LARP4B, LARP7) and heterogeneous nuclear ribonucleoproteins (hnRNPA3, hnRNPF, hnRNPH1, hnRNPU, hnRNPUL1) emerged as potential ITAFs to facilitate circRNA translation.

**Figure 3.**
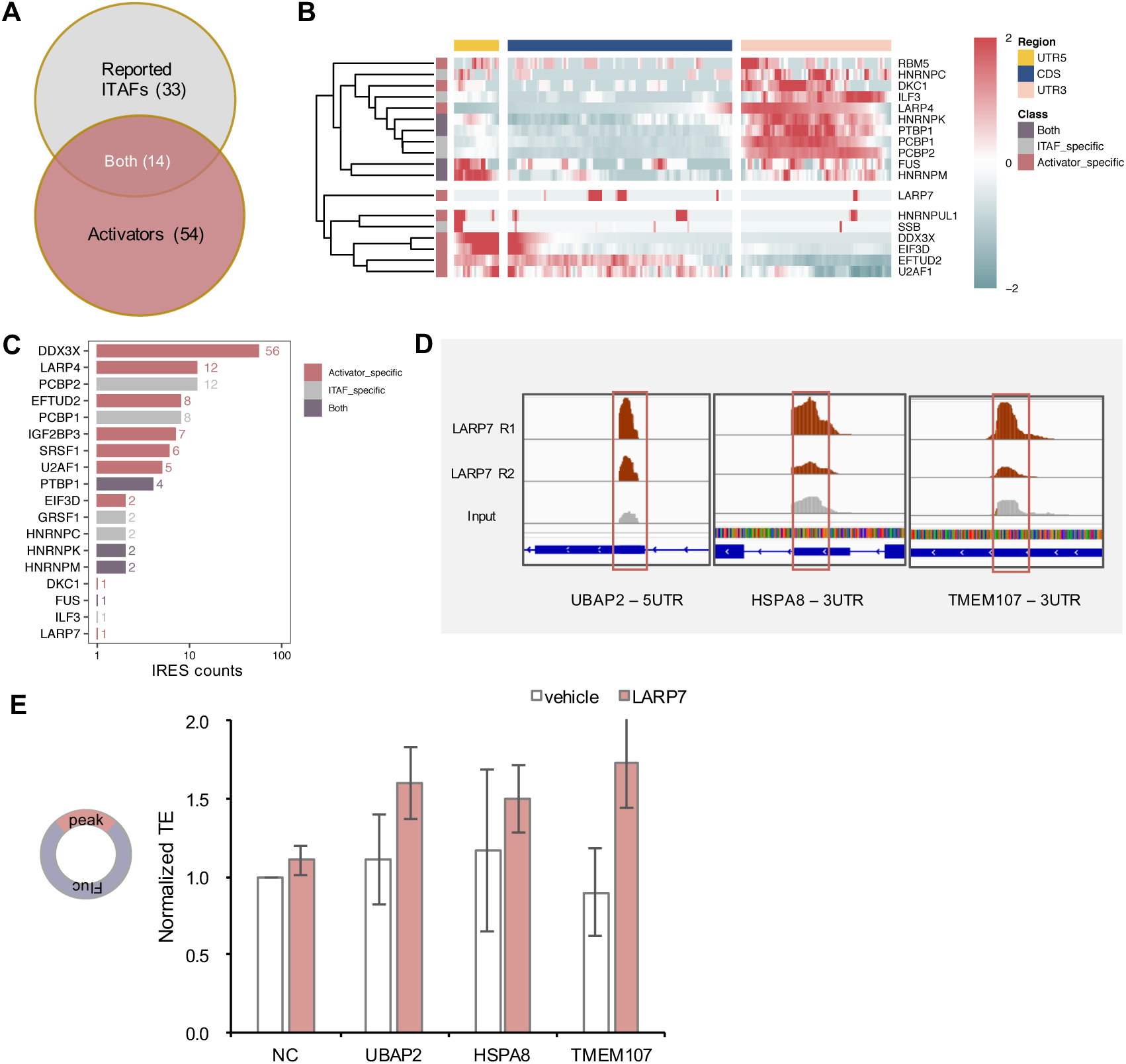
Activators can enhance circRNA translation through binding to endogenous RNA targets. (A) Venn diagram showing the overlap between identified activators and reported ITAFs. A total of 14 proteins were shared between the two groups, and there were 54 proteins unique to activators and 33 unique to reported ITAFs. (B) Peak distribution of reported ITAFs and identified activators using eCLIP data in HepG2 cell line. The yellow, dark blue and pink block represent UTR5, CDS and UTR3, respectively. (C) Counts of IRESs containing eCLIP peaks of different RBPs (within 150nt window at upstream and downstream of the IRES) in HepG2 cells. (D) Diagrams for the binding peaks of LARP7 along the transcripts of UBAP2, HSPA8 and TMEM107 in HepG2 cell line. (E) TE evaluation of circular luciferase reporter containing three eCLIP binding peaks of LARP7 (in the mRNA of UBAP2, HSPA8 and TMEM107), which were co-expressed with vehicle and pcDNA3.1- LARP7 in HepG2 cell line separately (mean ± SD, in triplicate transfections). NC was a designed non-targeting RNA control for LARP7.

To gain insight into how these newly identified translation activators may interact with endogenous mRNA, we analyzed the RNA-binding pattern of both identified activators and reported ITAFs using enhanced cross-linking immunoprecipitation (eCLIP) data generated by the ENCODE consortium in HepG2 and K562 cells (Van Nostrand et al., 2020). We first employed peak density as a metric to categorize RBPs by determining their binding preferences across the 5’ untranslated region (5’UTR), CDS and 3’ untranslated region (3’UTR) region of annotated transcripts. The majority of reported ITAFs and identified activators predominantly bound to the 5’UTR and 3’UTR regions, with the exception of DDX3X, eIF3D, EFTUD2 and U2AF1, which also exhibited binding peaks within the CDS region, in addition to 5’UTR in HepG2 cells (Fig 3B). The binding pattern was slightly different in K562 cell line, likely attributable to the influences of cell specificity and variations in eCLIP data quality (Supplementary Figure S4B).

To further explore the potential involvement of these translation activators in the IRES-dependent translation, we examined the correlation between eCLIP peaks of different activators and the known IRES regions using 693 human IRES sequences from IRESbase (Zhao et al., 2020). We measured the numbers of IRESs that contain nearby eCLIP peaks (within a 150 nt window) of different RBPs. Among these activators, DDX3X, LARP4 were ranked as the top two RBPs with overlapped IRES counts (Fig 3C). The DDX3X was reported to function as ITAF to promote IRES-dependent translation (Su et al., 2018). However the potential roles of La-related proteins as ITAFs to regulate cap-independent translation have not been carefully invested (Küspert et al., 2015).

We next used LARP7 as an example to further study its potential involvement in cap-independent translation. The eCLIP analysis in HepG2 cells was conducted to identify LARP7 binding peaks (methods), and we selected three distinct binding peaks within the 5’UTR of *Ubiquitin Associated Protein 2* (*UBAP2), Heat shock 70 kDa protein 8 (HSPA8)* and *Transmembrane Protein 107 (TMEM107)* genes for subsequent validation (Fig 3D). These endogenous LARP7 binding sites were then inserted between the start and stop codon of cFluc backbone. The resulting reporter was co-transfected with expression vector of FLAG tagged LARP7 to measure their effect on TE, and the FLAG tag plasmid was used as a vehicle control. As a result, the incorporation of LARP7 with cFluc containing peak sequences from *UBAP2*, *HSPA8* and *TMEM107* all exhibited an increased TE relative to the vehicle (Fig 3E). Our finding elucidated the potential role of LARP7 in modulating translation of circRNAs, implying that LARP7 may function as a putative ITAF *via* binding to its cellular RNA targets.

### Functional modules in the newly identified translation activators

Many RBPs have modular configuration, with the RNA binding domains to specifically recognize their targets and the functional domains to manipulate RNA metabolism (Lunde et al., 2007; Pereira et al., 2017). Notably, specific functional domains or unstructured fragments of RBPs can mimic the regulatory effects of the intact protein on regulating alternative splicing (Mao et al., 2018; Schmok et al., 2024), back-splicing (Qi et al., 2021) and translation (Reynaud et al., 2023; Villa and Fraser, 2024) when directed to their RNA targets. Building on these insights, we aimed to identify the particular domains or fragments of the activators that serve as functional modules to govern the translation of circRNAs.

We initially analyzed the annotated domains of the identified translation activators (Supplementary Figure S5A), and found many contain RNA helicase domains in addition to the canonical RNA-binding domains (like RRM or KH domain). However, the majority of the translation activators do not have well-defined domains that may serve as the functional modules. Guided by the annotated domains or regions from the UniProt database, we selected several representative activators and test the activity of their putative functional truncations using the tethering assay (Figure 4A). We first sought to examine activators with well-characterized domains, including helicase domain containing proteins (DDX21, eIF4A1 and SKIV2L2) and RRM containing proteins (FUS, RBM28 and LARP7). For translation activators without well-defined domains, such as eIF3D, eIF5B and SRRM1, we took intrinsically disordered regions (IDRs) into considerations. Additionally, we aimed to discern the translation regulatory patterns for NMD3, which contains a specific region essential for the nuclear export of the 60S ribosomal subunit (Lee et al., 2016; Trotta et al., 2003). hnRNPUL1 and PES1 were also selected in our study due to their high TE values (> 2) in our tethering assay. In total, the N-terminal and C-terminal truncations of 12 distinct translation activators were fused to the PUF domain, utilizing the identical backbone and conditions as our previously established PUF-RBPs library. These constructs were co-transfected with the cFluc reporter into the HEK 293T cell line to evaluate their TE values independently.

**Figure 4.**
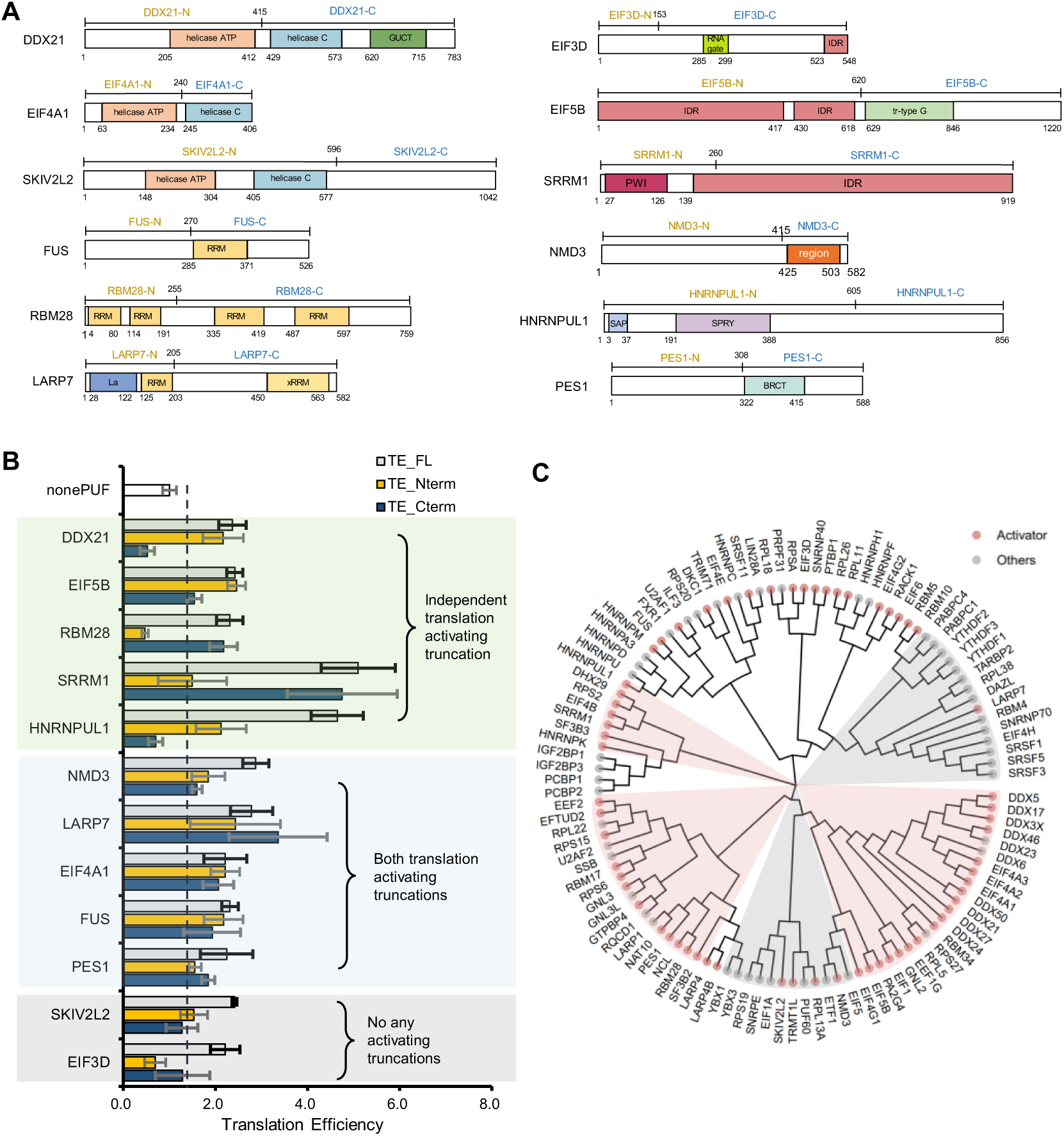
Domain analysis of identified activators reveals sequence similarity. (A) Diagram of domain configurations of the tested activators. The split position between N-terminal (yellow) and C-terminal (dark blue) fragments was marked. All tested truncations were shown. (B) Bar plot of TE values obtained from the tethering of full length protein (gray), N-terminal truncations (yellow) and C-terminal truncations (dark blue) of the 12 activators in HEK 293T (mean ± SD, in triplicate transfections). The dashed line shows the TE cutoff of 1.3. (C) Multiple amino acid sequence alignment of 110 RBPs performed using ClustalW, followed by clustering with MEGA X (neighbor-joining method). Red dots represent activators (TE ≥ 1.3) and gray dots represent others (TE < 1.3) in the screen.

We observed that, for some activators (including DDX21, eIF5B, RBM28, SRRM1), one of their N-terminal or C-terminal fragments exhibited comparable activity to the full-length protein. In particular, DDX21-Nterm (containing Helicase ATP), eIF5B-Nterm (containing 2 IDRs), RBM28-Cterm (containing 2 RRMs), and SRRM1-Cterm (containing one IDR) could recapitulate nearly all the translation regulatory activities of the intact protein (Figure 4B, Supplementary Figure S5B). On the other hand, the N-terminal fragment of hnRNPUL1 displayed approximately half of the activity in promoting TE compared to its full-length protein, whereas the hnRNPUL1-Cterm show no activity in increasing TE. In addition, for some translation activators (including NMD3, LARP7, eIF4A1, FUS, and PES1), both the N-terminal and C-terminal truncations were found to enhance circRNA translation. The activity of these truncations was generally not higher than the full-length proteins, implying that there are probably no inhibitory domains in these translation activators. Finally, for SKIV2L2 and eIF3D, none of the truncations can activate the circRNA translation like the full-length version (Figure 4B).

Interestingly, we found that the identity of annotated domains from these translation activators could not reliably predict their activity. For example, both eIF4A1 and SKIV2L2 contain a RNA helicase domain with two annotated subdomains (ATP-binding and C-terminal subdomains) (Figure 4A). However both subdomains showed translation promoting activity comparable to the full length eIF4A1, whereas the N-terminal truncation of SKIV2L2 containing the entire RNA helicase domain did not activate translation as judged by the same assay (Figure 4B). This result suggested that additional information besides the annotated domains may contribute to the translation promoting activities of RBPs. Considering these observations, we conducted a comprehensive amino acid sequences alignment from 110 RBPs and examined the association between the sequence compositions and their activities. Our analysis revealed a distinct clustering of the activators *vs.* the other proteins based on their sequences (Figure 4C), suggesting that the translation regulatory functions of these proteins may be closely associated with their amino acid compositions.

### Develop a machine learning model to analyze the translation activators of circRNAs

Our finding that the RBPs with similar activity can be clustered by their sequences (Figure 4C) implied that the sequence composition may be indicative to its function. In supporting of this notion, a recent study also reported that the activity of *trans*-acting splicing factors primarily relies on their sequence compositions (Mao et al., 2018). Therefore, we constructed a machine learning model to test if the activity of translation activators can be predicted by their amino acid sequences (Figure 2B). Specifically, we developed a Sparse Partial Least Square Discriminant Analysis (sPLS-DA) model using the 110 RBPs in our screen as the training set to learn their activity in regulating translation from the amino acid sequences. We first computed the frequencies of mono-peptide, di-peptide and tri-peptide across all the full length proteins, generating a 110 x 8420 input matrix (Figure 5A). The sparse features were subsequently filtered out using the sparsity cutoff at 0.95, resulting in 300 features that accounted for over 30% of the overall contribution (see methods). We used a 7-fold cross-validation to assess the model’s performance. Our model enabled a clear classification of the 110 RBPs into two distinct categories (activator *vs.* others), revealing the predictive power of short amino acid peptides in translation regulatory activity (Figure 5B). The average predictive accuracy for activators was above 70% as judged by various criteria for model performance (Figure 5C).

**Figure 5.**
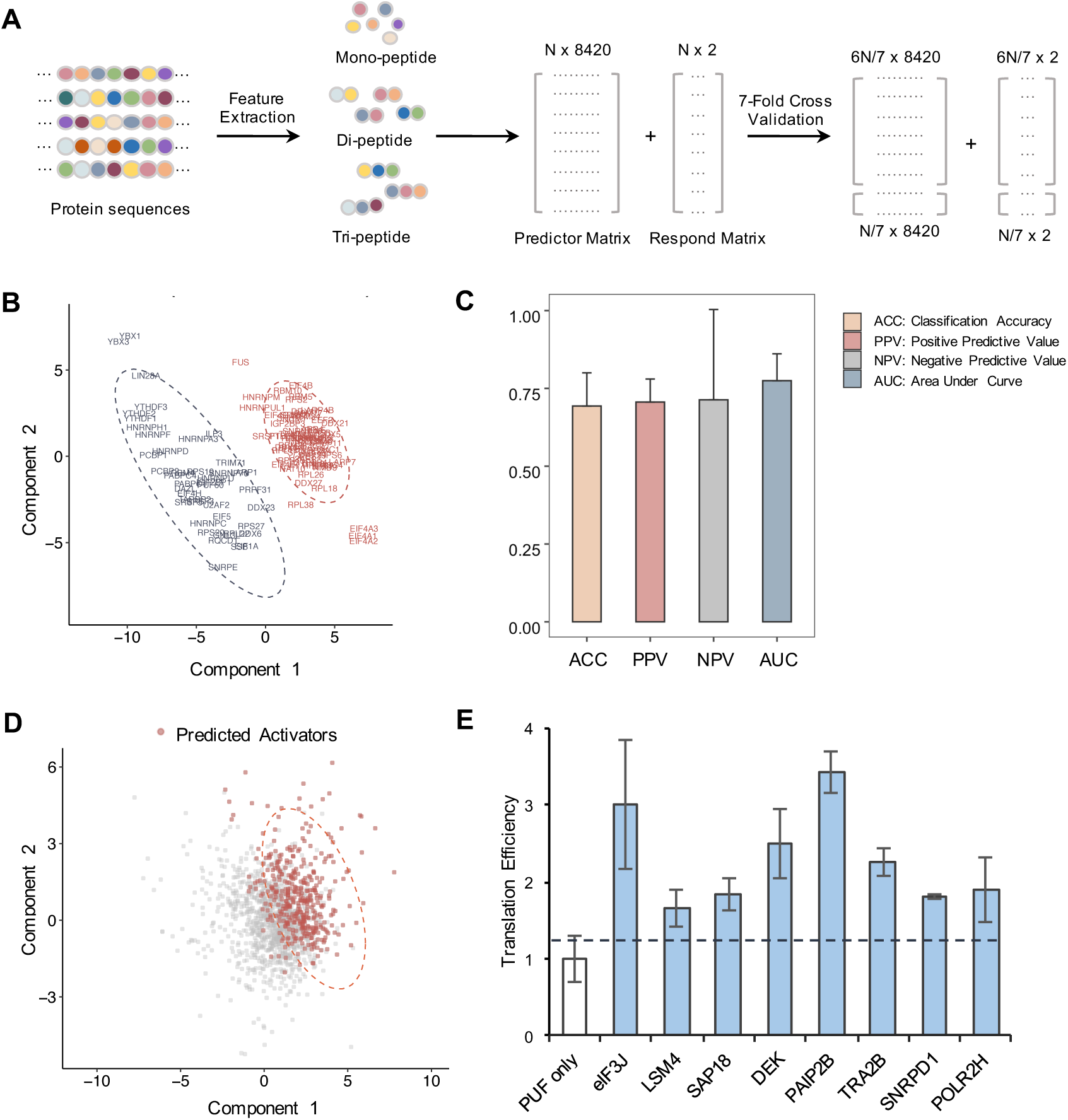
Prediction of novel activators using sPLS-DA model. (A) Workflow of the predictive model using sparse partial least-squares discriminant analysis (sPLS-DA). (B) Classification of the translation regulatory activities for all experimentally tested RBPs using sPLS-DA. The scatterplot shows the first two components in the principal component assay (PCA) using 300 selected features. Red fonts represent activators, and grey fonts represent others. The dashed line represents the 0.95 confidence ellipse for each category of RBPs. (C) The sPLS-DA prediction performance was evaluated using 7-fold cross-validation. The prediction accuracy (ACC), positive predictive value (PPV), negative predictive value (NPV), and area under the curve (AUC) were shown here (mean ± SD, flold k = 7, repeat n = 5). (D) Prediction of 1537 reported RBPs (excluding the 110 previous RBPs) for their regulatory activities in promoting circrRNA translation. Red fonts represent RBPs predicted as novel activators, and grey fonts represent others. The dashed line represents the 0.90 confidence ellipse for predicted activators. (E) Experimental evaluation of the ML-predicted activators in HEK 293T cells. The experimental details were the same as in Figure 2 (mean ± SD, in triplicate transfections). The dashed line shows the TE cutoff of 1.3.

We also examined the contribution of different sequence features to the model prediction, and found that the enrichment of K, L residues and IL, EK, PK dipeptides that may determine their activity as translation activators (Supplementary Figure S6A). We further applied this sPLS-DA model to the 1537 reported human RBPs in the UniProt database (Gerstberger et al., 2014) and predicted their translation promoting activity. Using a strict cutoff (probability ≥ 0.9), 398 proteins were predicted as translation activators (Figure 5D) (Table S3). The newly predicted activators also included some of unappreciated proteins in the eIF complexes, such as eIF3J that is a loosely associated subunit of human eIF3 complex (Zhou et al., 2008).

We next verified the reliability of our model by selecting eight candidates predicted as translation activators and testing their activities with the circRNA reporter system (Supplementary Figure S6B). Among the selected candidates, the oncoprotein DEK has recently been reported to participate in ribosome associated pathways through its interaction with DDX21 and RPL7a (Smith et al., 2018), whereas the PAIP2B was reported to inhibit translation of canonical mRNAs by displacing poly-A binding proteins (PABP) from the poly(A) tail (Berlanga et al., 2006). We found that all the candidates tested could increase circRNA translation efficiency compared to PUF only control (Figure 5E), demonstrating the accuracy of sPLS-DA model in predicting potential RBPs with translation regulatory activities. This ML model enabled artificial engineering of translation activators, which can be more useful in applying to circRNA translation control due to a shorter length (Supplementary Figure S6C). More importantly, these findings also provided new perspectives on how the sequences of RBPs may contribute to their translation regulatory activities, which may be applied in developing programmable RBPs to regulate IRES-mediated translation.

### The translation activators can function as ITAFs to promote IRES-mediated translation of circRNAs

The viral IRESs have been widely used in driving translation of circRNA (Chen et al., 2023; Wang and Wang, 2015; Wesselhoeft et al., 2018), which serve as the platform technology for a new generation of mRNA therapy (Wei et al., 2023). These IRESs were generally classified into four groups, three of which were reported to require the assistance of additional protein factors as ITAFs (Kieft, 2008; Yang and Wang, 2019). The group III and IV IRESs require the assistance of cellular auxiliary factors, including the eIFs and ITAFs, for efficient ribosome assembly, and the group II IRESs require only a subset of eIFs and some ITAFs for translation initiation (e.g. HCV IRES) (López-Ulloa et al., 2022; Yang and Wang, 2019). However, the group I IRESs are able to directly recruit ribosome for translation initiation without assistance of additional factors (Yang and Wang, 2019). We postulate that some of the newly identified translation activators may be used as ITAFs to promote IRES-dependent translation of circRNAs. To test this possibility, we designed circRNA reporters containing different IRES elements to examine if the recruitment of these factors may show ITAF-like activity. In total four IRES sequences were selected to represent the group II to IV viral IRESs, including the IRESs from the coxsackievirus B3 (CVB3, group IV), encephalomyocarditis virus (EMCV, group III), hepacivirus C (HCV, group II) and classical swine fever virus (CSFV, group II) (Fig 6A).

**Figure 6.**
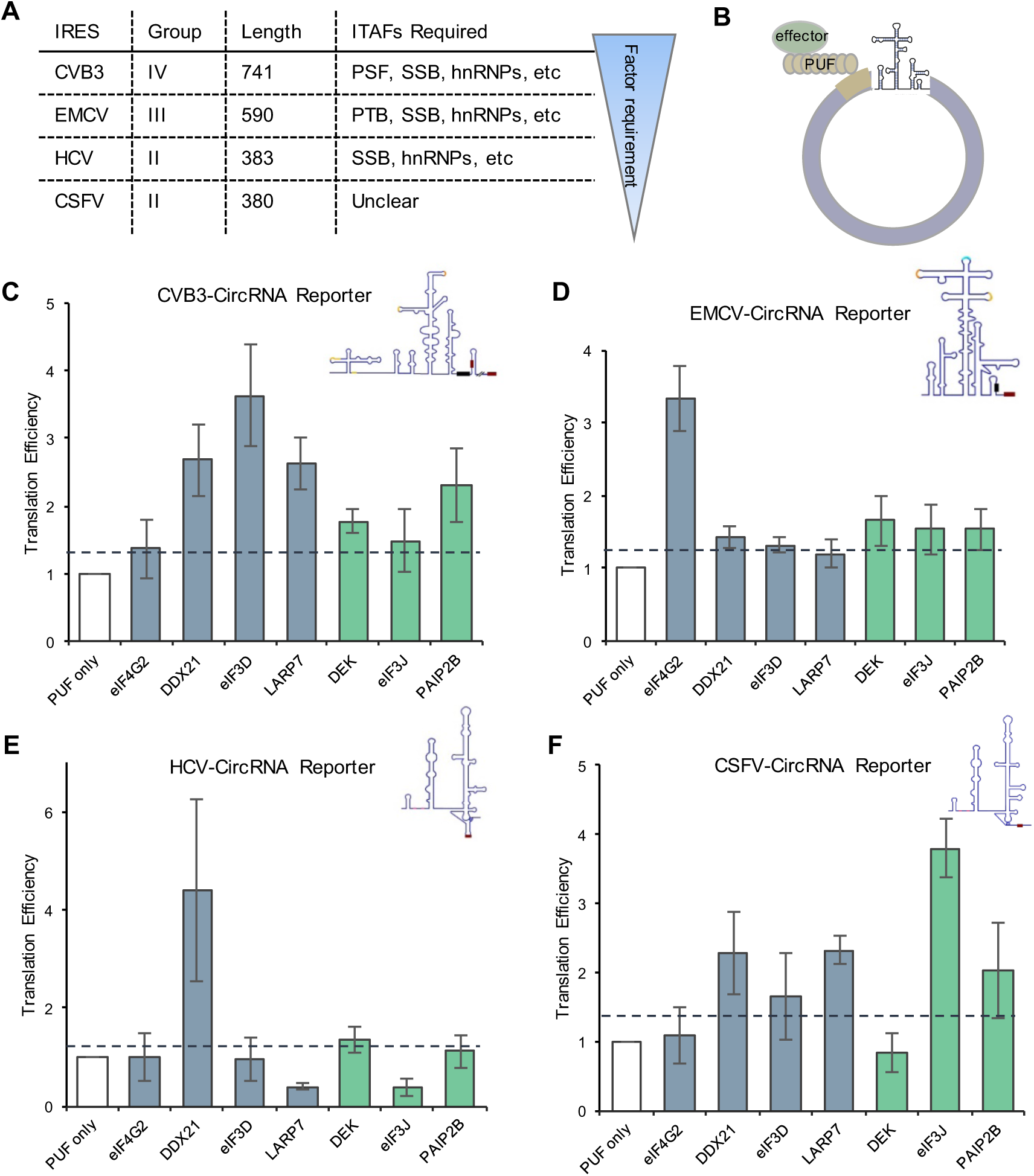
The newly identified activators can promote IRES-mediated circRNA translation. (A) Overview of four representative IRESs from three class: CVB3, EMCV, HCV, CSFV. (B) Diagram of the IRES-mediated circRNA translation reporter. Two tandem PUF binding sites were inserted between IRES sequence and stop codon of cFluc. (C-F) Bar plot displaying TE values of novel activators using dual circular luciferase reporter containing CVB3 (C), EMCV (D), HCV (E) and CSFV (F) respectively, as determined by the assay in Fig 1B (mean ± SD, in triplicate transfections). Experimentally validated translation activators from the initial screen were shown in gray blue, and RBPs from ML prediction were shown in green. The dashed line shows the TE cutoff of 1.3.

Each IRES sequence was inserted into a circular luciferase reporter containing an upstream site for the tethering of PUF fusion proteins (Fig 6B), and these reporters were co-expressed with PUF-RBPs to measure their activity in promoting circRNA translation (using the same assays as Fig 1B). We chose representative translation activators originally identified in our screen (DDX21, eIF3D, LARP7) and the activators identified through ML prediction (DEK, eIF3J, PAIP2B). We found that different combinations of IRESs and the translation activators showed a range of translation enhancement effects. Compared to the PUF only control, all the RBPs tested have increased translation of cFluc containing the group IV IRES from CVB3, with eIF3D displayed the highest activity in translation activation (Fig 6C). For the cFluc translation mediated by EMCV IRES, the tethering of the eIF4G2 showed the highest activity in translation enhancement (Fig 6D), with others have marginally increased translation (judged by the cutoff of TE > 1.3). For the IRESs from HCV and CSFV (both from group II), we found very distinct pattern in the effects of different RBPs: only DDX21 and DEK exhibited significant increase of TE values for HCV IRESs (Fig 6E), whereas most of the RBPs tested demonstrated translation promoting activities for CSFV IRESs (Fig 6F). Interestingly, although eIF4G2 was originally employed as a positive control in this assay, we found it only promote translation for the IRESs from CVB3 and EMCV (group IV and III), again indicating a large variation of RBP functions in different IRES contexts. Collectively, these results suggest that novel translation activators identified from tethering screen or ML prediction may be used to enhance IRES-mediated circRNA translation, however their activities may depend on the combination with distinct IRESs.

## Discussion

Optimizing translation efficiency is crucial for leveraging circRNAs as a new generation for mRNA drugs. Previous studies have primarily focused on the design and refinement of *cis*-elements (e.g. IRES, UTRs) (Chen et al., 2021a; Chen et al., 2023; Fan et al., 2022), while investigations into the role of *trans*- acting regulatory proteins have been relatively limited although they play critical roles in circRNA translation (Deviatkin et al., 2023; Sinha et al., 2022; Yang et al., 2017). In this study, we systematically investigated potential RBPs that can enhance cap-independent circRNA translation by screening a RBP tethering library. To our best knowledge, this work represents the first comprehensive exploration and predictive identification of potential ITAFs for circRNA translation, significantly expanding the scope of regulatory factors in cap-independent translation.

We employed the PUF domain to tether different RBPs and measured their activity in promoting circRNA translation. Compared to the tethering system using MS2 coat protein, our strategy offers more flexibility due to the programmable RNA-binding specificity of PUF, as it can be engineered to recognize endogenous sequences without insertion of multiple MS2 hairpins into the target. Previously, engineered PUF domains have been combined with various effector domains to manipulate different RNA processing pathways (Wei and Wang, 2015), including mRNA splicing (Wang et al., 2009) and translation (Cooke et al., 2011). In addition, Cao et al. engineered a light-inducible system to activate translation using the PUF scaffold with eIF4E (Cao et al., 2014). Our work has produced various artificial PUF factors with different functional domains, which may be useful in specifically modulating circRNA translation. Further optimizations could adapt the PUF domain to directly target *cis*-elements within endogenous circRNAs without changing their sequences. Moreover, the PUF domain may be further modified by including additional repeats to recognize longer RNA sequences (e.g., 10-12 nt), which can effectively increase its specificity and reduce off-target effects for *in vivo* applications (Han et al., 2022; Zhao et al., 2018).

We found that 68 RBPs out of 110 RBPs tested demonstrated the capacity to enhance the translation efficiency of circRNAs (Fig 2E). This seemingly high positive rate is probably because we pre-selected RBPs that were found to bind ribosomes. While almost all known ITAFs were studied in the context of linear mRNA, our results provide evidence that a subset of RBPs can enhance cap-independent translation for both circRNA and linear mRNA (Figure S3A, B). One possible explanation is that these proteins may play multi-functional roles by recruiting ribosomes and core translation initiation factors in synergy with IRES elements under cellular stress. It’s also worth noting that, even for linear mRNAs, translation predominantly occurs in a circularized form *via* the head-to-tail RNA looping mediated by the interaction between cap-binding proteins and the eIF4G (Wells et al., 1998). Therefore, the study on circRNA translation may also shed light on the regulation of linear mRNA. Moreover, elucidating the precise mechanisms by which RBPs regulate translation, and how to distinguish the IRES and ITAF functions in circRNA *vs.* linear RNA, necessitate further exploration.

Our results also showed that certain fragments of the RBPs can promote circRNA translation when tethered to circRNAs (Fig 4B). This discovery has two important implications. First, it suggests that truncated proteins retain the ability to recruit translation machinery and start initiation for circRNAs. Second, our approach offers a safer and more practical way to control translation with engineered proteins in mammalian cells, as the overexpression of a full-length RBP might interfere certain cellular processes by binding to its endogenous targets. Interestingly, we found that N-terminal IDR fragment of eIF5B may have independent activities in promoting translation. This observation aligns with *in vitro* findings that the disordered N-terminal region of eIF5B can stimulate the IRES activities in endogenous human genes by facilitating the formation of cytoplasmic granules (Harris and Marr, 2023). Importantly, the investigations of the activation domains within RBPs have been more thoroughly explored in other critical gene regulation processes including transcription (Alerasool et al., 2022; Ravarani et al., 2018; Tycko et al., 2020) and alternative splicing (Mao et al., 2018; Schmok et al., 2024). Given the conceptual parallels in different regulations of RNA processing, we anticipate that future screenings will be created to identify additional translation activation domains.

We further developed a ML approach using amino acid sequences to directly predict RBP function in controlling translation. AI-based approaches like AlphaFold have made great advances in predicting protein functions from specific structures (Abramson et al., 2024; Jumper et al., 2021). However, predicting the functions of unstructured peptides remains challenging. Our approach differs from the conventional function predictions in that it predicts the activities in protein or protein domains without clearly defined structures. In fact, we found that some of the unstructured IDRs (e.g., SRRM1 and eIF5B) are indeed responsible for promoting noncanonical translation (Fig 4B). A limitation of our ML model is the risk of false positive predictions, and the validation was restricted to a few RBPs. Future studies should explore additional RBPs involved in circRNA translation regulation, which may also consider additional parameters in refining the model. Despite this limitation, our study offers the possibility to predict translation activators based on their sequence compositions, expanding the scope of translation regulatory RBPs and allowing for the engineering of novel activators.

In summary, we conducted a systematic investigation into the *trans*-acting factors that promote the cap-independent circRNA translation. In addition to expanding the current knowledge on regulation of circRNA translation, this study may provide values in the application of circRNAs as new platform technology in RNA drug development. The specific expression profile of these translation activators may be used in controlling IRES-mediated translation of circRNAs under different cellular contexts. For example, we may use the expression preference of translation activators in different cell types and their cognate binding sites to engineer IRESs that can be “switched on” in certain cells, which is especially valuable in designing circRNAs with T-cell specific translation for cancer therapy. We anticipate that this line of research will offer a viable strategy for manipulating circRNA translation, thereby enhancing their applications in RNA therapy.

## Methods

### Construction of circular luciferase reporter

To generate circular RNA reporter, firefly luciferase gene was split into two fragments in reversed order, consistent with the split cFluc plasmid (Fan et al., 2022). Complementary elements flanked by introns were included to facilitate back-splicing by base pairing (Chen et al., 2021b). Cognate binding sites of the PUF domain were inserted into the middle of reverted luciferase fragments flanking with restriction endonucleases (EcoRI, SpeI). The construct was verified by Sanger sequencing, and primer information were listed in the Supplementary Table 1.

### Construction of RBP tethering library

To generate the candidate RBPs library, we firstly utilized the pool of proteins directly interacting with ribosome large subunit (data from the Supplementary Table S3 of (Simsek et al., 2017)) as the starting library. We intersected the ribosome-interacting proteins in the RNase-independent and puromycin-independent sets (using the NPV ≥ 0.99 as cutoff), resulting in 414 potential proteins. We further compared these 414 proteins with annotated RBPs (Gerstberger et al., 2014) to obtain 261 ribosome associated RBPs. Through manual refinement based on the existing literatures, we further reduced the list of 110 RBPs as the final collection.

To generate PUF tethered constructs for full length RBPs and C-terminal/N-terminal fragments (*i.e.*, effector domains), we used primers containing suitable restriction sites (XbaI, NotI) and human cDNAs for amplification (SuperScript III, Invitrogen). The amplified DNA fragments were then cloned into pGL-PUF backbone, flanking the corresponding restriction endonucleases by homologous recombination (ClonExpress^@^ Entry One Step Cloning Kit, Vazyme). A glycine rich linker was inserted between effector protein and PUF to enhance the conformational flexibility and stable expression of the fusion proteins (Reddy Chichili et al., 2013). The sequences of all cDNA clones was verified, and all primers and relevant information were included in the Supplementary Table 1.

### Cell culture and transfection

HEK 293T cells were cultured in DMEM medium supplemented with 10% FBS. One day before transfection, HEK 293T cells were seeded into 24-well plate, and transfection density was controlled at 35%-45%. For each transfection, 1.1 μL Lipofectamine 3000 (Invitrogen) and 0.72 μg plasmids (comprising 120 ng of reporter gene plasmid and 600 ng of PUF-RBP plasmid) were mixed with 50 μL Opti-MEM in separate tubes according to the manufacturer’s instructions. After gentle mixing and incubation for 15 minutes at room temperature, the solutions were added to 24-well plate and incubated in a 37°C incubator. The medium was changed after 8-12 hours, and cells were harvested 36 hours post-transfection.

### Dual luciferase assay

Cells in each well of 24-well plate were lysed with 200 uL Passive Lysis Buffer (Progema) at room temperature for 30 minutes. After overnight frozen at −80°C, half of the lysed cells were used for luciferase activity detection and western blotting, while the remaining half was reserved for RNA extraction. For luciferase activity detection, 10 uL of cell lysate was transferred to a 384-well plate. In the first reaction, 20 uL LARII (Progema) was added to assess the activity of firefly luciferase, followed by 20 uL Stop & Glo for Renilla luciferase with a parallel signal quenching of luciferase signal. The final luciferase activity was measured as the ratio of firefly luciferase activity normalized by internal Renilla luciferase activity.

### RNA extraction, semi-quantitative RT-PCR and real-time PCR

Total RNA was extracted from harvested transfected cells using TRIZOL reagent. For semi-quantitative PCR, 1ug of total RNA was treated with gDNA Eraser to remove genomic DNA, and then reverse transcribed with PrimeScript RT Reagent Kit (Takara). For real time PCR, one-tenth of the RT product was used as template using AceQ qPCR SYBR Green Master Mix (Vazyme) on a Roche LC480 system, following the manufacturer’s instructions. The amount of circRNA and circRNA back-splicing efficiency were measured by RT-PCR using designed primers (listed in the Supplementary Table 1).

### Western blotting

The total cells were lysed in 1×SDS-PAGE loading buffer (a mixture of 5×SDS protein loading buffer (Beyotime), RIPA buffer and protease inhibitor), heated at 98°C for 10 min. The samples were separated by 4-20% SurePAGE™ Gel (GenScript) for 1 h at 140V and then transferred to PVDF membranes for 2 h at 75V. The following antibodies was used: Flag antibody (F1084, Sigma-Aldrich) is diluted by 1:2000; beta Actin antibody (66009-1-Ig, Proteintech) is diluted by 1:10000. The HRP-linked second antibodies were used by 1:2000 dilution. The blots were visualized using an enhanced chemiluminescence detection kit and followed by ChemiDoc Touch Imaging System (Image Lab, Bio-Rad).

### Analysis of meta-transcripts peak density of eCLIP-seq data

To analyze the distribution of RBP binding sites across different transcript regions, we first downloaded all RBP idr (irreproducible discovery rate) peak files (GENCODE version 19) from eCLIP-seq data on the ENCODE project website (https://www.encodeproject.org), covering 103 RBPs from HepG2 cell line and 120 RBPs from K562 cell line. Transcripts abundance in the cell lines was derived from ENCODE’s rRNA-depleted RNA-seq data (HepG2: ENCFF000FVT, ENCFF000FVU; K562: ENCFF000HJF, ENCFF000HJX). We filtered for highly expressed (TPM ≥ 1) transcripts to serve as representative transcripts. To prevent redundancy from multiple isoforms of the same gene, we selected the transcript with the highest expression level per gene, yielding 10,722 transcripts in HepG2 and 9,624 transcripts in K562 for subsequent analysis.

We further calculated peak density of 5’UTR, CDS, and 3’UTR among exon regions of each transcript. Then the three regions of each transcript were divided into bins and the average numbers of peaks for exonic genomic locus were calculated within each bin.

To evaluate the appropriate numbers of bins, we calculated the median lengths of the 5’UTR, CDS, and 3’UTR regions in transcripts with TPM > 10. Based on these medians, the transcripts in HepG2 were divided into 182 bins (20 in 5’UTR, 100 in CDS, and 67 in 3’UTR) and the transcripts in K562 were divided into 175 bins (20 in 5’UTR, 100 in CDS, and 55 in 3’UTR).

Finally, we normalized the peak coverage of each bin in each transcript by dividing the sum of coverage of all transcripts. The sum of the normalized coverages on all transcripts represented the distribution of specific RBP binding sites and we defined these values as the meta-transcripts densities. For visualization, we calculated the z-score value of the average density across bins for each RBP and represented using the R package pheatmap.

### Analysis of the overlapped eCLIP-seq peaks with IRES regions

To seek the translation regulation evidence *via* IRES elements, we calculated the overlaps between eCLIP peaks and IRES regions. We first downloaded the 693 human IRES sequences from IRESbase (Zhao et al., 2020) and then defined the upstream and downstream exonic 150nt around IRES as IRES regions. The overlapped sequences were extracted using in-house python script.

### Sparse Partial Least Squares Discriminant Analysis (sPLS-DA)

Existing supervised machine learning models, such as random forests and support vector machines, face limitations when processing feature matrices characterized by high multicollinearity and sparsity. To address this issue, we employed a supervised machine learning algorithm, sparse version of Partial Least Squares Regression (sPLS-DA) to classify the translation activators according to their activities.

The sPLS-DA algorithm incorporates Lasso regularization into the PLS loading vector during the singular value decomposition (SVD) process (Rohart et al., 2017), and allows for dimensionality reduction, feature extraction, and discriminant analysis of sparse matrices (Lê Cao et al., 2011; Mao et al., 2018). In our analysis, we structured the predictor dataset, X, structured as an *n* × *p* matrix, where *n* represents the number of observations and *p* denotes the number of variables. The response dataset, *Y*, was organized as an *n* × *q* matrix with *n* observations and *q* classes (where *q* = 2). This response dataset was recoded into a dummy matrix suitable for classification purposes in PLS. The PLS algorithm constructs a set of orthogonal components designed to maximize the sample covariance between the response and the linear combination of the predictor variables:

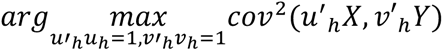

where *u_h_* and *v_h_* represent the left and right singular vectors for the SVD of *X^T^Y* respectively in each iteration or dimension *h* of the PLS (where *h* = 1 … *H*, *H* is the chosen number of dimensions). The original approach was based on SVD and the optimization problem of loading vectors when 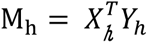 is:

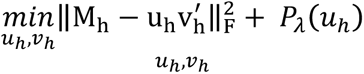

where *P_λ_*(*u_h_*) is the soft thresholding function that approximates Lasso penalty function.

In the discrimination step, the PLS model can be formulated as *Y* = *Xβ* + *E*, where *β* is the matrix of the regression coefficients and *E* is the residual matrix. The coefficient matrix *β* is calculated as *β* = *W*^∗^*V*^*T*^, where *V* is the matrix of right singular vectors derived from the SVD decomposition (columns *v_1_*, …, *v_H_*) and *W*^∗^ = *W*(*u^T^W*)^-1^. Here, *W* represents the matrix of regression coefficients from the regression of *X* on the latent variable *t_h_* = *v*′*_h_Y*, and *u* is the matrix of left singular vectors (columns *u_1_*,…, *u_H_*).

In the predictive step, a new set of samples is calculated as *Y_test_* = *X_test_β*, where *β* is the same matrix of the regression coefficients, *X_test_* is the predictor matrix for test samples. The classification of each test sample (each row in *Y_test_*) assigns the column index of the element with the highest predicted value within that row.

### sPLS-DA Model Construction

Based on data from the initial screening, we classified the 110 RBPs into two groups, activator and others, using a translation efficiency (TE) threshold of 1.3. To represent the amino acid composition of each full-length protein, we then extracted the frequencies of short-peptide features, which include mono-peptide, di-peptide, and tri-peptide (see Figure 5A). Features with high sparsity (> 0.95) were excluded from the analysis, and the remaining features were then input into the sPLS-DA model, which was implemented using the R package *mixOmics*. Model performance was evaluated through 7-fold cross-validation, conducted with the R package *crossval*. Short-peptide frequencies were calculated by in-house Python script. The figures were generated using ggplot2. The machine learning prediction values for the reported RBPs can be found in Supplementary Table 3.

## Data availability

The data in this study is available as the supplementary tables, and software code used in this study are available in https://github.com/rnasys/CITActivator.

## Acknowledgements

We thank members and alumni of the Wang lab, in particular F.X.J, L.J.F, W.H.H and M.M.W for advice and support. We thank Dr. Xiaowei Song for sharing a dozen of plasmids containing full length RBPs.

The work is funded by National Key Research and Development Program of China (MOST grant #2021YFA1300503 to ZW), National Natural Science Foundation of China (NSFC grant #32030064 and #32250013 to ZW), the Strategic Priority Research Program of Chinese Academy of Sciences (grant #XDB38040100 to ZW), Starry Night Science Fund at Shanghai Institute for Advanced Study of Zhejiang University (SN-ZJU-SIAS-009 to Z.W.).

## Author contributions

L.Q.Y. was primarily responsible for designing and executing experiments, part of data analysis. L.Q.Y. prepared the original manuscript and W.Z.F. revised the manuscript. W.S.Q. performed computational analysis in this work and wrote the manuscript. Y.Y.W carried out several of the experiments, under the supervision of L.Q.Y. and W.Z.F.. Y.Y. consulted the screening experiment in the project.

## Declaration of AI-assisted technologies in the writing process

During the preparation of this work, we used Chatgpt in order to optimize the grammar and wording. After using this tool, we reviewed and edited the content as needed and take full responsibility for the content of the publication.

## Reference

Abe, N., Matsumoto, K., Nishihara, M., Nakano, Y., Shibata, A., Maruyama, H., Shuto, S., Matsuda, A., Yoshida, M., Ito, Y., et al. (2015). Rolling Circle Translation of Circular RNA in Living Human Cells. Sci Rep 5, 16435.

Abramson, J., Adler, J., Dunger, J., Evans, R., Green, T., Pritzel, A., Ronneberger, O., Willmore, L., Ballard, A.J., Bambrick, J., et al. (2024). Accurate structure prediction of biomolecular interactions with AlphaFold 3. Nature 630, 493–500.

Alerasool, N., Leng, H., Lin, Z.-Y., Gingras, A.-C., and Taipale, M. (2022). Identification and functional characterization of transcriptional activators in human cells. Molecular cell 82, 677–695.e677.

Benoit Bouvrette, L.P., Wang, X., Boulais, J., Kong, J., Syed, Easin U., Blue, Steven M., Zhan, L., Olson, S., Stanton, R., Wei, X., et al. (2022). RBP Image Database: A resource for the systematic characterization of the subcellular distribution properties of human RNA binding proteins. Nucleic Acids Research 51, D1549–D1557.

Berlanga, J.J., Baass, A., and Sonenberg, N. (2006). Regulation of poly (A) binding protein function in translation: Characterization of the Paip2 homolog, Paip2B. RNA 12, 1556–1568.

Calviello, L., Mukherjee, N., Wyler, E., Zauber, H., Hirsekorn, A., Selbach, M., Landthaler, M., Obermayer, B., and Ohler, U. (2016). Detecting actively translated open reading frames in ribosome profiling data. Nature methods 13, 165–170.

Cao, J., Arha, M., Sudrik, C., Schaffer, D.V., and Kane, R.S. (2014). Bidirectional regulation of mRNA translation in mammalian cells by using PUF domains. Angew Chem Int Ed Engl 53, 4900–4904.

Carter, M.S., and Sarnow, P. (2000). Distinct mRNAs that encode La autoantigen are differentially expressed and contain internal ribosome entry sites. Journal of Biological Chemistry 275, 28301–28307.

Chang, J., Shin, M.-K., Park, J., Hwang, H.J., Locker, N., Ahn, J., Kim, D., Baek, D., Park, Y., Lee, Y., et al. (2023). An interaction between eIF4A3 and eIF3g drives the internal initiation of translation. Nucleic Acids Research 51, 10950–10969.

Chen, C.-K., Cheng, R., Demeter, J., Chen, J., Weingarten-Gabbay, S., Jiang, L., Snyder, M.P., Weissman, J.S., Segal, E., Jackson, P.K., et al. (2021a). Structured elements drive extensive circular RNA translation. Molecular Cell 81, 4300–4318.e4313.

Chen, C., Yang, Y., and Wang, Z. (2021b). Study of circular RNA translation using reporter systems in living cells. Methods 196, 113–120.

Chen, C.Y., and Sarnow, P. (1995). Initiation of protein synthesis by the eukaryotic translational apparatus on circular RNAs. Science 268, 415–417.

Chen, R., Wang, S.K., Belk, J.A., Amaya, L., Li, Z., Cardenas, A., Abe, B.T., Chen, C.- K., Wender, P.A., and Chang, H.Y. (2023). Engineering circular RNA for enhanced protein production. Nature biotechnology 41, 262–272.

Chen, T.-M., Lai, M.-C., Li, Y.-H., Chan, Y.-L., Wu, C.-H., Wang, Y.-M., Chien, C.-W., Huang, S.-Y., Sun, H.S., and Tsai, S.-J. (2019). hnRNPM induces translation switch under hypoxia to promote colon cancer development. EBioMedicine 41, 299–309.

Conn, V.M., Chinnaiyan, A.M., and Conn, S.J. (2024). Circular RNA in cancer. Nature Reviews Cancer 24, 597–613.

Cooke, A., Prigge, A., Opperman, L., and Wickens, M. (2011). Targeted translational regulation using the PUF protein family scaffold. Proc Natl Acad Sci U S A 108, 15870–15875.

Deviatkin, A.A., Simonov, R.A., Trutneva, K.A., Maznina, A.A., Soroka, A.B., Kogan, A.A., Feoktistova, S.G., Khavina, E.M., Mityaeva, O.N., and Volchkov, P.Y. (2023). Cap-Independent Circular mRNA Translation Efficiency. Vaccines (Basel) 11, 238.

Fan, X., Yang, Y., Chen, C., and Wang, Z. (2022). Pervasive translation of circular RNAs driven by short IRES-like elements. Nature Communications 13, 3751.

Feng, X.-Y., Zhu, S.-X., Pu, K.-J., Huang, H.-J., Chen, Y.-Q., and Wang, W.-T. (2023). New insight into circRNAs: characterization, strategies, and biomedical applications. Experimental Hematology & Oncology 12, 91.

Gerstberger, S., Hafner, M., and Tuschl, T. (2014). A census of human RNA-binding proteins. Nat Rev Genet 15, 829–845.

Godet, A.C., David, F., Hantelys, F., Tatin, F., Lacazette, E., Garmy-Susini, B., and Prats, A.C. (2019). IRES Trans-Acting Factors, Key Actors of the Stress Response. Int J Mol Sci 20.

Han, S., Sun, S., Li, P., Liu, Q., Zhang, Z., Dong, H., Sun, M., Wu, W., Wang, X., and Guo, H. (2020). Ribosomal protein L13 promotes IRES-driven translation of foot-and-mouth disease virus in a helicase DDX3-dependent manner. Journal of Virology 94, 10.1128/jvi.01679-01619.

Han, W., Huang, W., Wei, T., Ye, Y., Mao, M., and Wang, Z. (2022). Programmable RNA base editing with a single gRNA-free enzyme. Nucleic Acids Research 50, 9580–9595.

Harris, M.T., and Marr, M.T. (2023). The intrinsically disordered region of eIF5B stimulates IRES usage and nucleates biological granule formation. Cell reports 42.

Hinnebusch, A.G. (2014). The scanning mechanism of eukaryotic translation initiation. Annu Rev Biochem 83, 779–812.

Holdt, L.M., Kohlmaier, A., and Teupser, D. (2018). Circular RNAs as therapeutic agents and targets. Frontiers in physiology 9, 1262.

Jumper, J., Evans, R., Pritzel, A., Green, T., Figurnov, M., Ronneberger, O., Tunyasuvunakool, K., Bates, R., Žídek, A., and Potapenko, A. (2021). Highly accurate protein structure prediction with AlphaFold. Nature 596, 583–589.

Kieft, J.S. (2008). Viral IRES RNA structures and ribosome interactions. Trends in biochemical sciences 33, 274–283.

Kristensen, L.S., Andersen, M.S., Stagsted, L.V.W., Ebbesen, K.K., Hansen, T.B., and Kjems, J. (2019). The biogenesis, biology and characterization of circular RNAs. Nature Reviews Genetics 20, 675–691.

Kristensen, L.S., Jakobsen, T., Hager, H., and Kjems, J. (2022). The emerging roles of circRNAs in cancer and oncology. Nature Reviews Clinical Oncology 19, 188–206.

Küspert, M., Murakawa, Y., Schäffler, K., Vanselow, J.T., Wolf, E., Juranek, S., Schlosser, A., Landthaler, M., and Fischer, U. (2015). LARP4B is an AU-rich sequence associated factor that promotes mRNA accumulation and translation. RNA 21, 1294–1305.

Kwan, T., and Thompson, S.R. (2019). Noncanonical Translation Initiation in Eukaryotes. Cold Spring Harb Perspect Biol 11.

Lê Cao, K.-A., Boitard, S., and Besse, P. (2011). Sparse PLS discriminant analysis: biologically relevant feature selection and graphical displays for multiclass problems. BMC bioinformatics 12, 1–17.

Lee, A.S., Kranzusch, P.J., Doudna, J.A., and Cate, J.H. (2016). eIF3d is an mRNA cap-binding protein that is required for specialized translation initiation. Nature 536, 96–99.

Legnini, I., Di Timoteo, G., Rossi, F., Morlando, M., Briganti, F., Sthandier, O., Fatica, A., Santini, T., Andronache, A., and Wade, M. (2017). Circ-ZNF609 is a circular RNA that can be translated and functions in myogenesis. Molecular cell 66, 22–37. e29.

Liu, C.-X., and Chen, L.-L. (2022). Circular RNAs: Characterization, cellular roles, and applications. Cell 185, 2016–2034.

López-Ulloa, B., Fuentes, Y., Pizarro-Ortega, M.S., and López-Lastra, M. (2022). RNA-Binding Proteins as Regulators of Internal Initiation of Viral mRNA Translation. Viruses 14, 188.

Lunde, B.M., Moore, C., and Varani, G. (2007). RNA-binding proteins: modular design for efficient function. Nature reviews Molecular cell biology 8, 479–490.

Mao, M., Hu, Y., Yang, Y., Qian, Y., Wei, H., Fan, W., Yang, Y., Li, X., and Wang, Z. (2018). Modeling and Predicting the Activities of Trans-Acting Splicing Factors with Machine Learning. Cell Systems 7, 510–520 e514.

Marash, L., Liberman, N., Henis-Korenblit, S., Sivan, G., Reem, E., Elroy-Stein, O., and Kimchi, A. (2008). DAP5 promotes cap-independent translation of Bcl-2 and CDK1 to facilitate cell survival during mitosis. Molecular cell 30, 447–459.

Niu, D., Wu, Y., and Lian, J. (2023). Circular RNA vaccine in disease prevention and treatment. Signal Transduction and Targeted Therapy 8, 341.

Pamudurti, N.R., Bartok, O., Jens, M., Ashwal-Fluss, R., Stottmeister, C., Ruhe, L., Hanan, M., Wyler, E., Perez-Hernandez, D., Ramberger, E., et al. (2017). Translation of CircRNAs. Molecular cell 66, 9–21 e27.

Patop, I.L., Wust, S., and Kadener, S. (2019). Past, present, and future of circRNAs. EMBO J 38, e100836.

Pereira, B., Billaud, M., and Almeida, R. (2017). RNA-binding proteins in cancer: old players and new actors. Trends in cancer 3, 506–528.

Perenkov, A.D., Sergeeva, A.D., Vedunova, M.V., and Krysko, D.V. (2023). In Vitro Transcribed RNA-Based Platform Vaccines: Past, Present, and Future. 11, 1600.

Petkovic, S., and Müller, S. (2015). RNA circularization strategies in vivo and in vitro. Nucleic Acids Research 43, 2454–2465.

Qi, Y., Han, W., Chen, D., Zhao, J., Bai, L., Huang, F., Dai, Z., Li, G., Chen, C., Zhang, W., et al. (2021). Engineering circular RNA regulators to specifically promote circular RNA production. Theranostics 11, 7322–7336.

Ravarani, C.N., Erkina, T.Y., De Baets, G., Dudman, D.C., Erkine, A.M., and Babu, M.M. (2018). High-throughput discovery of functional disordered regions: investigation of transactivation domains. Molecular Systems Biology 14, e8190.

Reddy Chichili, V.P., Kumar, V., and Sivaraman, J. (2013). Linkers in the structural biology of protein–protein interactions. Protein science 22, 153–167.

Reynaud, K., McGeachy, A.M., Noble, D., Meacham, Z.A., and Ingolia, N.T. (2023). Surveying the global landscape of post-transcriptional regulators. Nature Structural & Molecular Biology 30, 740–752.

Riggs, C.L., Kedersha, N., Ivanov, P., and Anderson, P. (2020). Mammalian stress granules and P bodies at a glance. Journal of Cell Science 133, jcs242487.

Rohart, F., Gautier, B., Singh, A., and Lê Cao, K.-A. (2017). mixOmics: An R package for ‘omics feature selection and multiple data integration. PLoS computational biology 13, e1005752.

Schmok, J.C., Jain, M., Street, L.A., Tankka, A.T., Schafer, D., Her, H.-L., Elmsaouri, S., Gosztyla, M.L., Boyle, E.A., Jagannatha, P., et al. (2024). Large-scale evaluation of the ability of RNA-binding proteins to activate exon inclusion. Nature biotechnology 42, 1429–1441.

Simsek, D., Tiu, G.C., Flynn, R.A., Byeon, G.W., Leppek, K., Xu, A.F., Chang, H.Y., and Barna, M. (2017). The Mammalian Ribo-interactome Reveals Ribosome Functional Diversity and Heterogeneity. Cell 169, 1051–1065 e1018.

Sinha, T., Panigrahi, C., Das, D., and Chandra Panda, A. (2022). Circular RNA translation, a path to hidden proteome. Wiley Interdisciplinary Reviews: RNA 13, e1685.

Smith, E.A., Krumpelbeck, E.F., Jegga, A.G., Prell, M., Matrka, M.M., Kappes, F., Greis, K.D., Ali, A.M., Meetei, A.R., and Wells, S.I. (2018). The nuclear DEK interactome supports multi-functionality. Proteins: Structure, Function, Bioinformatics 86, 88–97.

Su, Y.-S., Tsai, A.-H., Ho, Y.-F., Huang, S.-Y., Liu, Y.-C., and Hwang, L.-H. (2018). Stimulation of the internal ribosome entry site (IRES)-dependent translation of enterovirus 71 by DDX3X RNA helicase and viral 2A and 3C proteases. Frontiers in Microbiology 9, 382467.

Trotta, C.R., Lund, E., Kahan, L., Johnson, A.W., and Dahlberg, J.E. (2003). Coordinated nuclear export of 60S ribosomal subunits and NMD3 in vertebrates. The EMBO Journal 22, 2841–2851.

Tycko, J., DelRosso, N., Hess, G.T., Aradhana, Banerjee, A., Mukund, A., Van, M.V., Ego, B.K., Yao, D., Spees, K., et al. (2020). High-throughput discovery and characterization of human transcriptional effectors. Cell 183, 2020–2035.e2016.

Van Nostrand, E.L., Freese, P., Pratt, G.A., Wang, X., Wei, X., Xiao, R., Blue, S.M., Chen, J.Y., Cody, N.A.L., Dominguez, D., et al. (2020). A large-scale binding and functional map of human RNA-binding proteins. Nature 583, 711–719.

Villa, N., and Fraser, C.S. (2024). Human eukaryotic initiation factor 4G directly binds the 40S ribosomal subunit to promote efficient translation. Journal of Biological Chemistry 300.

Wang, Y., Cheong, C.-G., Tanaka Hall, T.M., and Wang, Z. (2009). Engineering splicing factors with designed specificities. Nature methods 6, 825–830.

Wang, Y., and Wang, Z. (2015). Efficient backsplicing produces translatable circular mRNAs. RNA 21, 172–179.

Wei, H.-H., Zheng, L., and Wang, Z. (2023). mRNA therapeutics: New vaccination and beyond. Fundamental Research 3, 749–759.

Wei, H., and Wang, Z. (2015). Engineering RNA-binding proteins with diverse activities. Wiley Interdisciplinary Reviews: RNA 6, 597–613.

Wells, S.E., Hillner, P.E., Vale, R.D., and Sachs, A.B. (1998). Circularization of mRNA by eukaryotic translation initiation factors. Molecular cell 2, 135–140.

Wesselhoeft, R.A., Kowalski, P.S., and Anderson, D.G. (2018). Engineering circular RNA for potent and stable translation in eukaryotic cells. Nat Commun 9, 2629.

Wesselhoeft, R.A., Kowalski, P.S., Parker-Hale, F.C., Huang, Y., Bisaria, N., and Anderson, D.G. (2019). RNA Circularization Diminishes Immunogenicity and Can Extend Translation Duration. Molecular Cell 74, 508–520.

Yang, Y., Fan, X., Mao, M., Song, X., Wu, P., Zhang, Y., Jin, Y., Yang, Y., Chen, L.L., Wang, Y., et al. (2017). Extensive translation of circular RNAs driven by N(6)- methyladenosine. Cell Research 27, 626–641.

Yang, Y., Gao, X., Zhang, M., Yan, S., Sun, C., Xiao, F., Huang, N., Yang, X., Zhao, K., and Zhou, H. (2018). Novel role of FBXW7 circular RNA in repressing glioma tumorigenesis. JNCI: Journal of the National Cancer Institute 110, 304–315.

Yang, Y., and Wang, Z. (2019). IRES-mediated cap-independent translation, a path leading to hidden proteome. Journal of Molecular Cell Biology 11, 911–919.

Zhao, J., Li, Y., Wang, C., Zhang, H., Zhang, H., Jiang, B., Guo, X., and Song, X. (2020). IRESbase: a comprehensive database of experimentally validated internal ribosome entry sites. Genomics, Proteomics & Bioinformatics 18, 129–139.

Zhao, Y.-Y., Mao, M.-W., Zhang, W.-J., Wang, J., Li, H.-T., Yang, Y., Wang, Z., and Wu, J.-W. (2018). Expanding RNA binding specificity and affinity of engineered PUF domains. Nucleic acids research 46, 4771–4782.

Zhou, M., Sandercock, A.M., Fraser, C.S., Ridlova, G., Stephens, E., Schenauer, M.R., Yokoi-Fong, T., Barsky, D., Leary, J.A., and Hershey, J.W. (2008). Mass spectrometry reveals modularity and a complete subunit interaction map of the eukaryotic translation factor eIF3. Proceedings of the National Academy of Sciences 105, 18139–18144.

