## Supplementary Figures for "Systematic screen of RNA binding proteins that enhance circular RNA translation"

Supplementary Figure S1 (related to Figure 2): Functional annotation and expression of RBPs used in the tethering screen.

Supplementary Figure S2 (related to Figure 2): The effect of the selected RBPs on back-splicing efficiency.

Supplementary Figure S3 (related to Figure 2): Additional tests on identified translation activators.

Supplementary Figure S4 (related to Figure 3): Computational analyses of the known ITAFs and the identified translation activators.

Supplementary Figure S5 (related to Figure 4): Domain analyses on the 68 identified translation activators.

Supplementary Figure S6 (related to Figure 5): Sequence features of the translation activators predicted by our ML method.

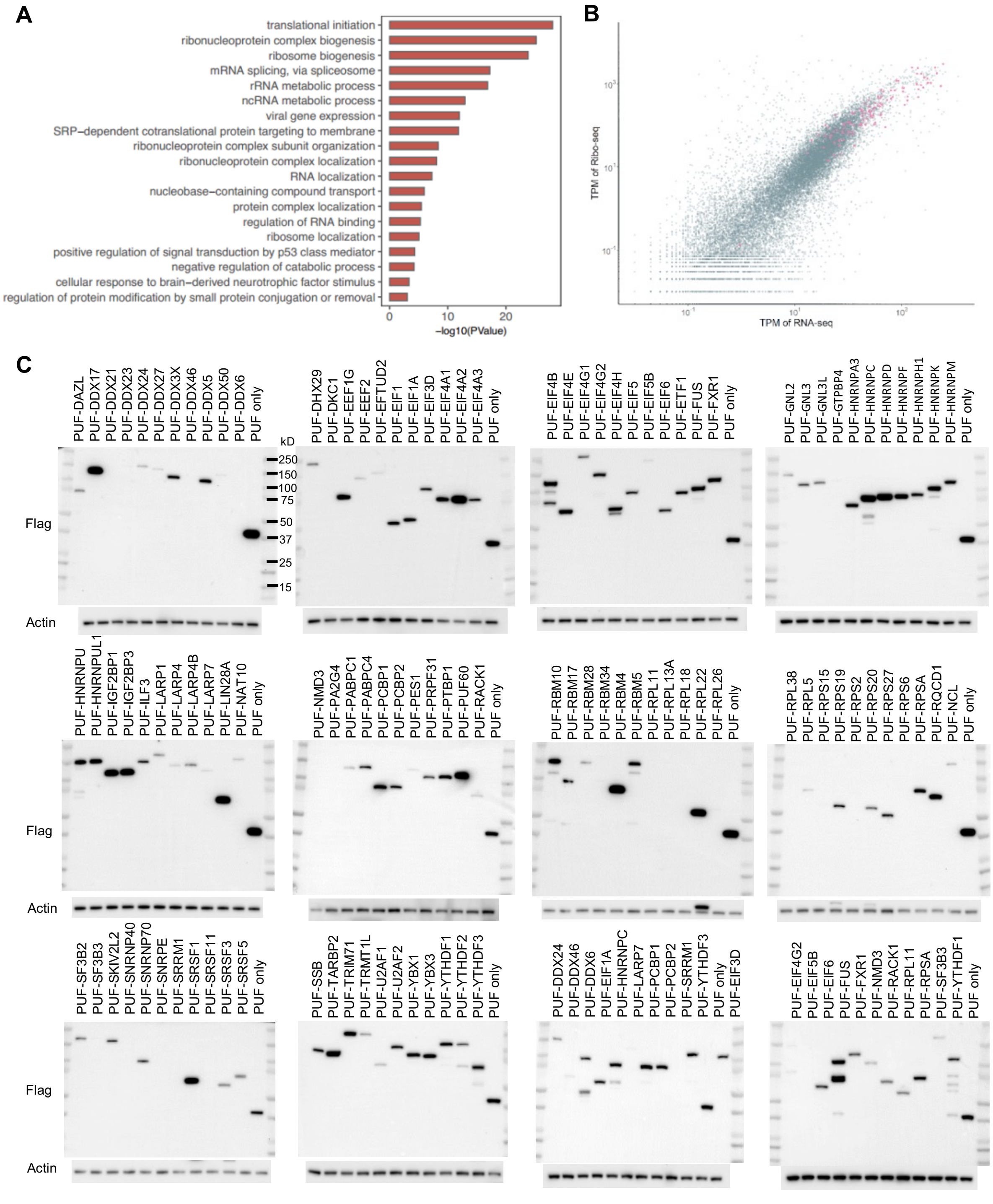


**Figure S1. Functional annotation and expression of RBPs used in the tethering screen.**

(A) GO analysis of 110 RBPs that are known to interact with ribosomes based on previous studies.

(B) Correlation between the RNA levels and the translation of all genes in HEK 293T cells. The transcript per million (TPM) values from RNA-seq and TPM of Ribo-seq data were represented by a dot plot, and the 110 RBPs were marked by purple dots.
(C) Confirmation of the expression for the 110 PUF-RBP fusion proteins using western blotting analysis.

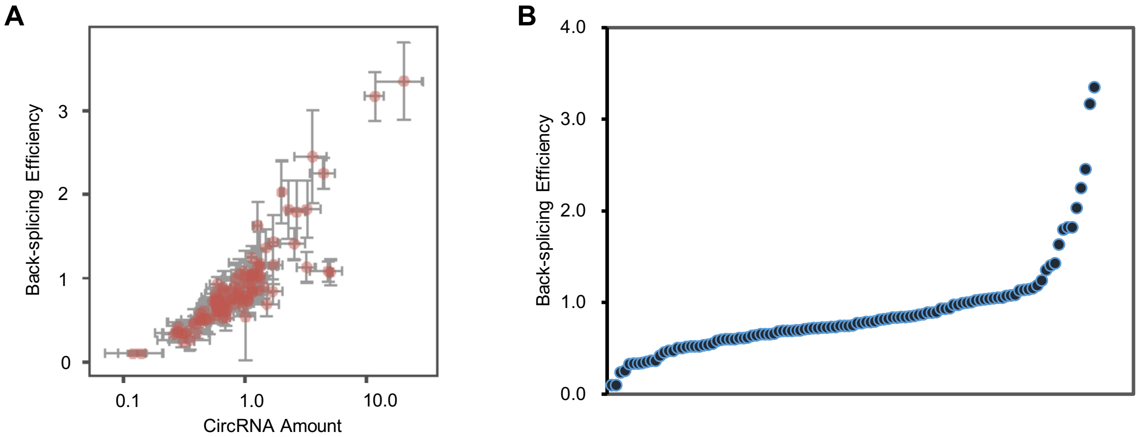


**Figure S2. The effect of the selected RBPs on back-splicing efficiency.**

(A) The scatter plot showing the correlation between back-splicing efficiency and circRNA amount, both values normalized to PUF-only.

(B) The scatter plot showing the distribution of back-splicing efficiency for all tested RBPs.

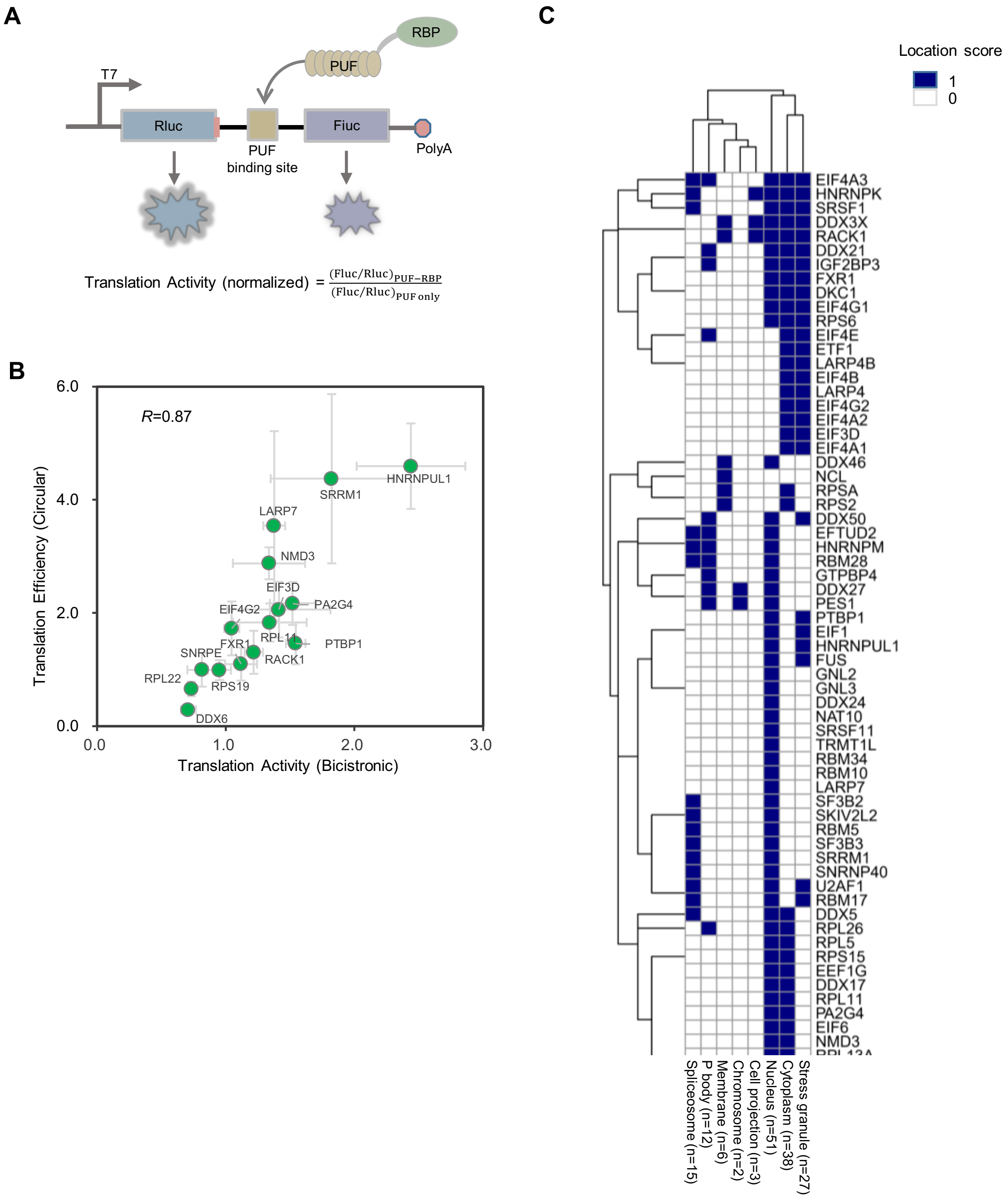


**Figure S3. Additional tests on identified translation activators.**

(A) Diagram of the standard bicistronic reporter. Two tandem PUF binding sites were inserted 8 bp before the Fluc start codon, and tested PUF-RBPs were co-transfected with the reporter in HEK 293T cells. 36 hours post-transfection, cells were lysed, and luminescence was measured using a microplate reader to calculate translation activity. PUF only was used as control vector.

(B) Scatter plot of translational efficiency for tested RBPs in circular RNA and their translation activities in the bicistronic reporter (mean ± SD, in triplicate transfections). *R* represents the Pearson correlation coefficient.

(C) Clustering of location scores for 68 translation activators, including their presence in the nucleus (n=51), cytoplasm (n=38), stress granules (n=27), P-bodies (n=12), etc.

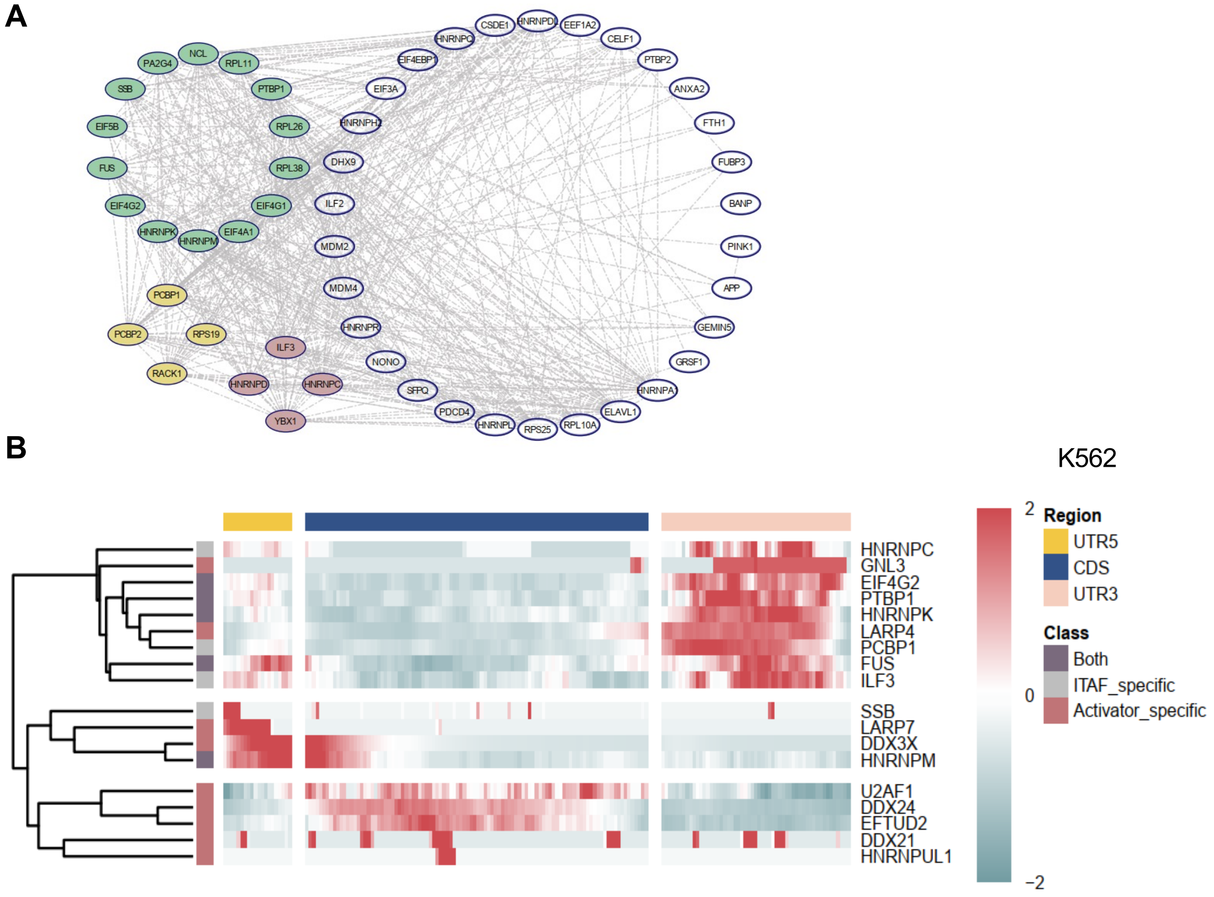


**Figure S4. Computational analyses of the known ITAFs and the identified translation activators.**

(A) Protein-protein interaction network of 55 reported ITAFs using STRING database and CytoScape software, which is comparable to that of the identified translation activators (main Fig. 2). Gemin5, ELAVL1, PTBP1 were absent in the network for limited interactions with other proteins as determined by STRING database.

(B) Peak distribution plot of reported ITAFs and identified activators using eCLIP data in the K562 cell line.

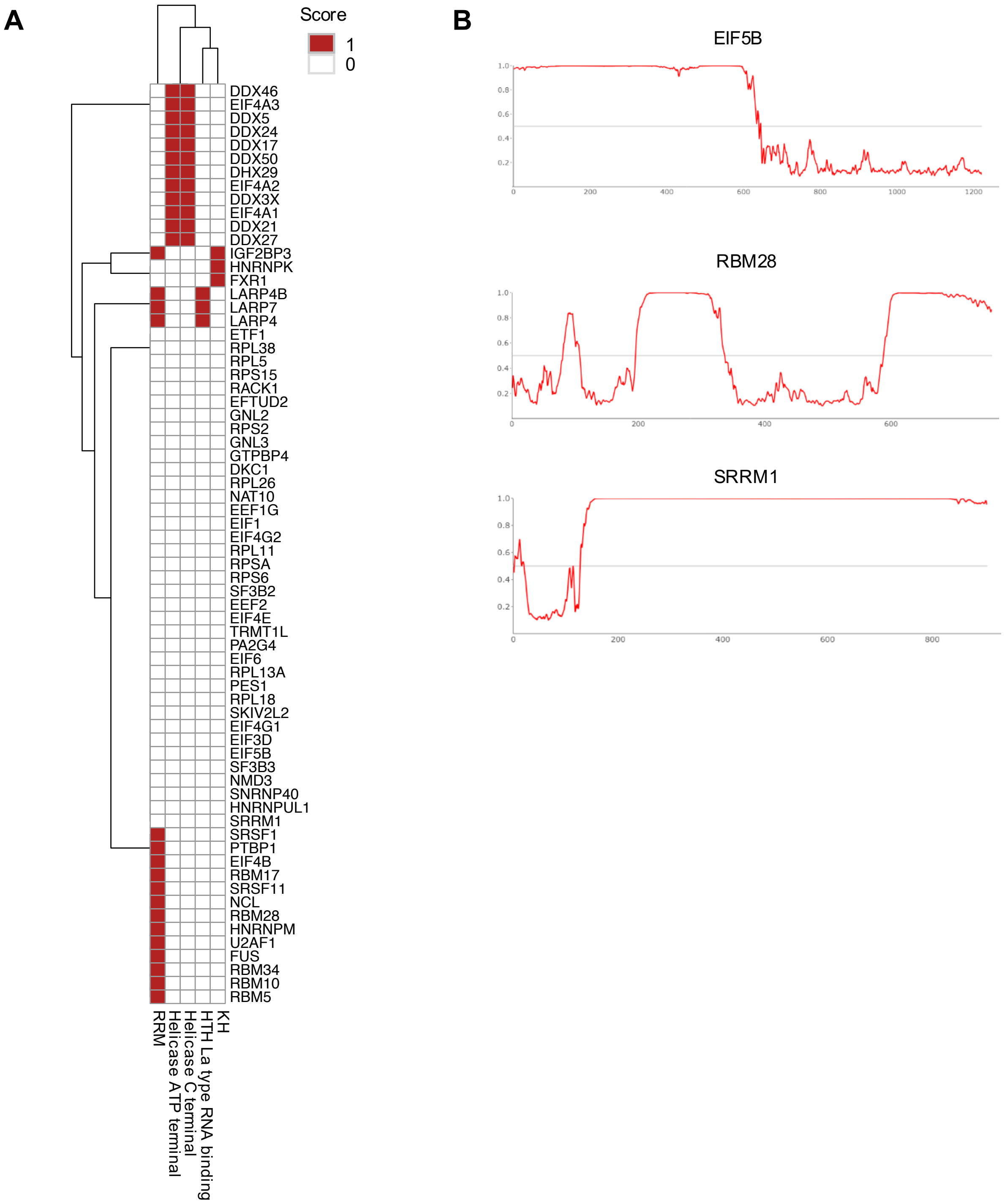


**Figure S5. Domain analyses on the 68 identified translation activators.**

(A) Clustering of structured domains (annotated in Uniprot database) in 68 identified activators.

(B) IDR prediction plots of EIF5B, RBM28 and SRRM1 within their full length proteins using AIUPred database (Nucleic Acids Research (2024): gkae385.).

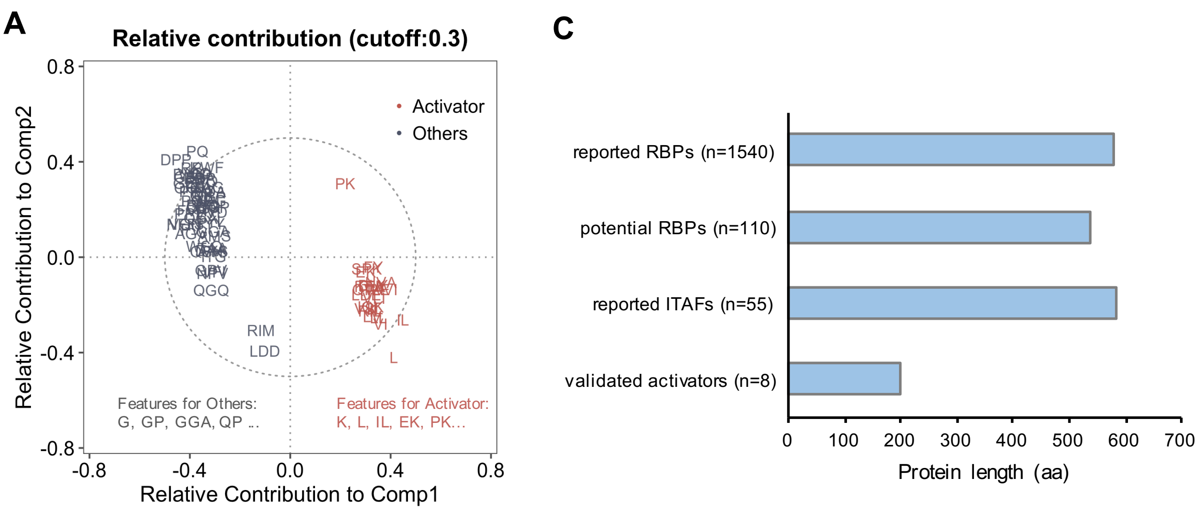


**B**

Predicted ML score > 0.5

100aa < Protein length < 400aa

| | **eIF3J** | **LSM4** | **SAP18** | **DEK** | **PAIP2B** | **TRA2B** | **SNRPD1** | **POLR2H** |
|---|---|---|---|---|---|---|---|---|
| ML Score | 0.56 | 0.66 | 0.67 | 0.77 | 0.80 | 0.93 | 0.95 | 1.32 |
| Protein length (aa) | 258 | 139 | 153 | 375 | 123 | 288 | 119 | 150 |

**Figure S6. Sequence features of the translation activations predicted by our ML method.**

(A) Relative contributions of various sequence features in the predictive model, calculated by the loading matrix of sPLS-DA model. Red fonts represent predictive features for translation activators and gray fonts represent others.

(B) Selection of RBPs for validation from the perspective of predicted probability (score > 0.5) and protein length (from 100 to 400 amino acids). The predicted ML scores and protein length for the 8 validated RBPs were listed in the table.

(C) Average protein length of reported RBPs (n = 1540), potential RBPs (n = 110), reported ITAFs (n = 55) and validated activators (n = 8).
